# Sparsely Phosphorylable Residues, Tuned by Agonist Chemotype, Shapes GPCR Signaling

**DOI:** 10.64898/2026.10.06.756976

**Authors:** Madhurjya Protim Borah, Sarfraz Ahmed, Mahek Agrawal, Neeraj Soni, Punita Kumari, Mithu Baidya

## Abstract

GPCR signaling diversity is often attributed to an intracellular phosphorylation “barcode,” in which ligand-dependent modification of multiple receptor phosphoresidues encodes selective engagement of downstream transducers. However, whether such an extensive phosphorylation of the receptor is universally required remains unclear. Here, we show that the melanocortin-4 receptor (MC4R), a central regulator of energy homeostasis, achieves remarkably diverse and pathway-selective signaling through a sparse set of strategically positioned regulatory phosphoresidues. Systematic study of these phosphoresidues within helix 8 and the C-terminal tail revealed distinct, spatially segregated contributions to receptor function. Substitution of helix 8 residues S306 disrupted β-arrestin2 recruitment, early endosomal trafficking, and ERK1/2 activation, while visibly altering Gαs–cAMP signaling in all the receptor variants. In contrast, distal C-terminal substitutions had little effect on β-arrestin2 coupling and trafficking, revealing that these regions are not functionally redundant. Surprisingly, signaling through Gα_q/11_-and Gα_12/13_-linked pathways remained largely constrained across receptor variants, further demonstrating pathway-specific control by these phosphoresidues. Notably, ligand chemotype reshaped the functional consequences of these substitutions: setmelanotide selectively restored or enhanced signaling through pathways that were compromised by specific receptor mutations. Together, our findings argue that GPCR signaling diversity does not necessarily require an extensive phosphorylation barcode. Instead, MC4R uses a minimal number of spatially organized phosphoresidue that can differentially control transducer engagement and downstream signaling, while ligand chemistry determines how this regulatory landscape is accessed.

## 1. Introduction

G protein-coupled receptors (GPCRs) are among the most versatile signaling proteins in the cell, translating an extraordinary diversity of extracellular cues, from photons and ions to peptides and small molecules, into precisely regulated intracellular responses. Despite this diversity, the core molecular mechanism of GPCR signaling is remarkably conserved (*1*). Agonist binding reshapes the receptor to engage intracellular transducers, while receptor phosphorylation and other intracellular regulatory events help determine the duration, location, and nature of the resulting signal (*2*). Thus, much of the functional diversity observed across receptors originates from which receptor conformations and transducers are engaged at the receptor itself in response to different ligands. One mechanism proposed to regulate this choice is the GPCR “phospho-code,” in which agonist-dependent phosphorylation of intracellular receptor regions allows the receptor to model a multitude of different conformations that influences β-arrestin engagement and downstream signaling (*3*, *4*). The original barcode model envisioned an extensive pattern of phosphorylation across receptor tails, but subsequent structural and biochemical studies have progressively refined this view, showing how strategically positioned phosphorylation sites stabilize functionally distinct β-arrestin states (*5*, *6*). This raises an important question: when a receptor contains only a limited set of intracellular regulatory residues, can individual residues exert disproportionate and pathway-selective effects on signaling?

The melanocortin-4 receptor (MC4R) provides an especially compelling system in which to address this question. MC4R is a central regulator of energy homeostasis, and pathogenic variants in MC4R represent the most common monogenic cause of severe human obesity (*7*). Although MC4R has traditionally been studied through its canonical Gα_s_-cAMP pathway, it is now clear that the receptor engages a broader signaling repertoire that includes Gα_q/11_, Gα_12/13_ (*8–11*), β-arrestin2, and ERK/MAPK (*12–15*). Yet the intracellular regulatory architecture of MC4R is relatively sparse compared with receptors classically associated with elaborate phosphorylation barcodes. Its helix 8 (H8) and C-terminal regions contain a limited number of serine and threonine residues, several of which have been implicated in receptor regulation (*16*). This creates a striking paradox: how can a receptor with such minimal intracellular regulatory residues generate such diverse signaling outputs?

Interestingly, human genetic studies provide an important clue. Naturally occurring MC4R variants can produce highly selective changes in receptor signaling, and variation in β-arrestin-associated signaling has been linked to metabolic phenotypes independently of canonical Gα_s_-cAMP output (*17*). These observations challenge the notion that a disease-associated MC4R variant simply causes a global loss of receptor function across all downstream signaling pathways. Instead, they suggest that disruption of individual intracellular residues can selectively alter receptor coupling to specific transducers, such that impairment of one signaling pathway can occur without equivalent disruption of others. This raises a fundamental question: how can a receptor with such a sparse intracellular regulatory architecture generate and differentially control such a broad signaling repertoire? A further layer of complexity comes from the ligand. Different agonists can stabilize distinct receptor conformations, thereby altering transducer engagement and signaling output (*18–20*). Thus, the functional consequence of an MC4R mutation may depend not only on the affected residue but also on the ligand that occupies the receptor. This is particularly relevant to MC4R, where chemically diverse agonists, including melanocortin peptides, small molecules, and the clinically approved agonist setmelanotide that displays distinct signaling profiles (*21*, *22*). Whether ligand structure can systematically reshape the signaling consequences of individual H8 and C-terminal mutations, and whether such remodeling is shared across transducers or restricted to particular mutant–transducer combinations, remains unknown.

Here, we address this question by systematically interrogating selected H8 and C-terminal serine/threonine residues across Gα_s_, β-arrestin2, receptor trafficking, ERK, Gα_q/11_, and Gα_12/13_ signaling. By combining these mutations with a chemically diverse agonist panel, we ask how a limited set of intracellular residues can encode pathway-selective signaling and how ligand structure modulates these effects. Importantly, inclusion of S306, which has not been established as a phosphorylation site, allows us to determine whether this residue contributes to the regulation of MC4R signaling despite its unestablished phosphorylation status.

## 2. Result

### 2.1. Identification of candidate regulatory phosphorylation sites within the H8 and C-terminal tail of MC4R

The intracellular helix 8 (H8) and C-terminal tail are key regulatory elements of Class A GPCRs, where their sequence and conformational properties contribute to receptor coupling, phosphorylation-dependent regulation, β-arrestin recruitment, desensitization, and trafficking. In several GPCRs, receptor activation is accompanied by conformational rearrangements of H8 (*23–27*) exposing regulatory serine and threonine residues that might influence GRK-mediated phosphorylation while also facilitating interactions with intracellular signaling and trafficking proteins including βarrs. These observations suggest that H8, together with the C-terminal tail, functions as a dynamic signaling platform that fine-tunes receptor activity. Consistent with other Class A GPCRs, MC4R contains a short amphipathic H8 located immediately downstream of the highly conserved NPxxY motif at the cytoplasmic end of transmembrane helix 7. H8 is anchored to the plasma membrane through palmitoylation of C318 and is followed by the intracellular C-terminal tail (Fig. 1). Given the established regulatory role of these intracellular regions in GPCR signaling, we first sought to identify candidate phosphorylation sites within H8 and the C-terminal tail of MC4R. Sequence analysis revealed two potential phospho-acceptor residues within H8 (S306 and T312) and two within the C-terminal tail (S329 and S330) (Fig. 1A). Mining of the PhosphoSitePlus database together with published studies identified T312 and the S329/S330 cluster as previously reported or putative phosphorylation sites, whereas S306 has not been described as a phosphosite (*16*). Interestingly, multiple sequence alignment demonstrated that S306 is highly conserved across vertebrate MC4R orthologs, despite the absence of experimental evidence for its phosphorylation (Fig. 1B). Such strong evolutionary conservation suggests that S306 may serve an important structural and/or regulatory function. Thus, based on their localization within established GPCR regulatory domains, existing biochemical studies (*16*), and evolutionary conservation, we selected S306, T312, S329, and S330 for systematic functional characterization (Fig. 1C) to define their contributions to MC4R activation, βarr recruitment, receptor trafficking, and downstream signaling.

**Figure 1.**
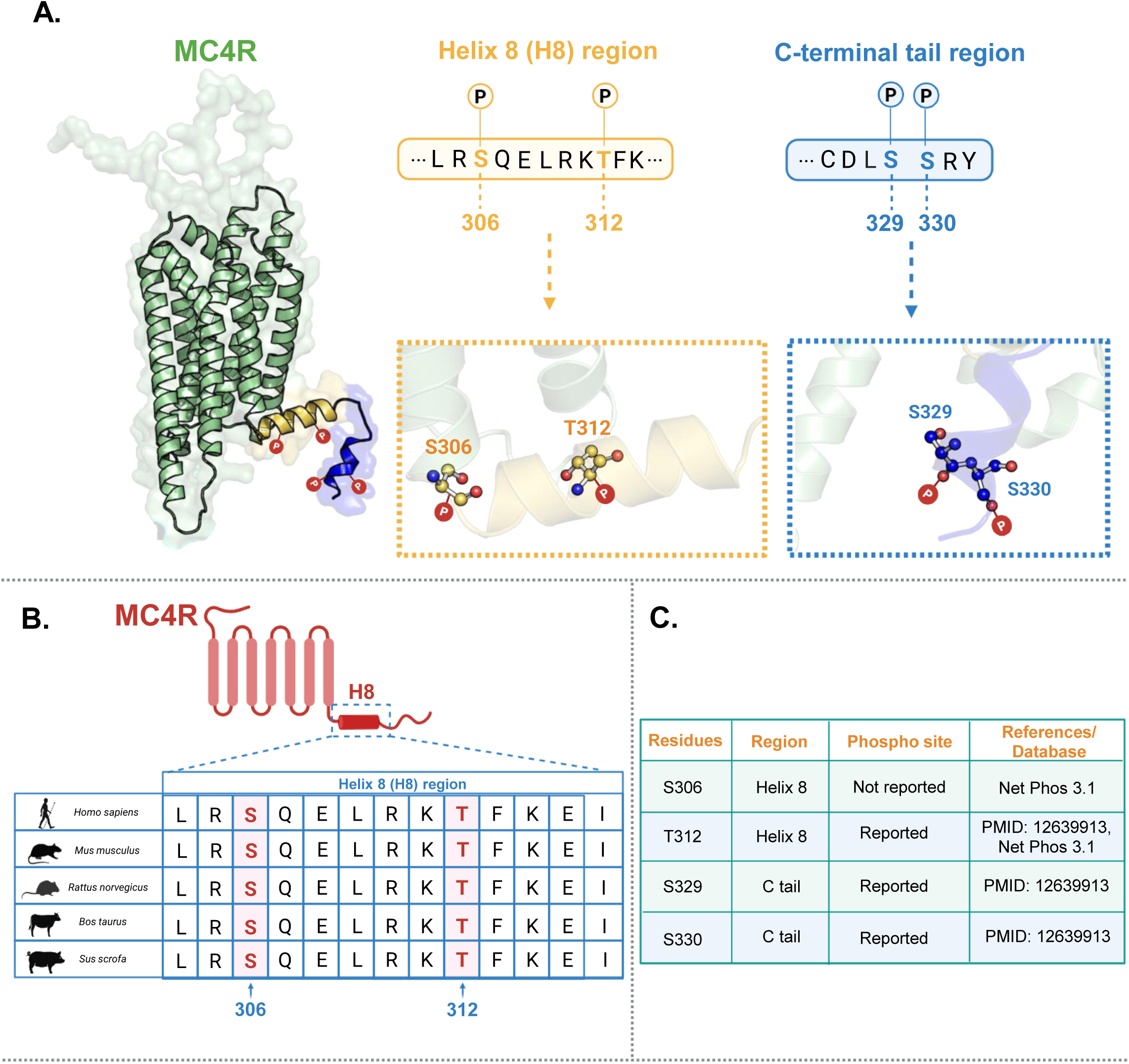
Structural location, sequence conservation, and reported phosphorylation status of selected serine/threonine residues in Helix 8 and C-tail of MC4R. **A.** Structural representation of MC4R highlighting four conserved serine/threonine residues. S306 and T312 are located in the helix 8 (H8) region (Yellow helix) and S329 and S330 are located in the C-terminal tail (Blue helix). Zoomed inset shows the ball and stick orientation of each residue and phosphorylation sites/potential phosphorylation sites are indicated by the addition of “P”. **B.** Multiple sequence alignment of the MC4R H8 region across different species shows the conservation of S306 and T312. **C.** Table showing summary of the reported/unreported phosphorylation status of S306, T312, S329, and S330. S306 has not been reported as phosphorylated, whereas T312, S329, and S330 have been previously reported as phosphorylation sites.

### 2.2 Functional characterization of regulatory residues within helix 8 and the C-terminal tail of MC4R

Having identified candidate regulatory S/T residues within helix 8 (S306 and T312) and the C-terminal tail (S329 and S330) of MC4R, we next sought to define the contribution of each residue to receptor activation and signaling. We first generated individual alanine substitutions to abolish the hydroxyl side chain and prevent phosphorylation at these positions (Fig. 2A). Before assessing receptor function, we first determined whether these substitutions affected receptor biosynthesis or plasma membrane localization using whole-cell ELISA performed in HEK293T cells. Our results demonstrated that all mutants were expressed at the cell surface at levels comparable to MC4R^WT^ (Fig. S1), indicating that the mutations did not compromise receptor folding, maturation, or forward trafficking. These findings thus validate the mutant receptors for subsequent functional analyses. To understand the contribution of these individual residues on receptor activation, we measured Gα_s_-mediated signaling using a luminescence-based GloSensor cAMP assay following α-MSH stimulation. Although all mutants reached the plasma membrane efficiently, they exhibited distinct signaling phenotypes (Fig. 2, B to E). Mutations within both H8 and the C-terminal tail reduced agonist-stimulated signaling relative to MC4R^WT^ resulting in a decrease in maximal response (*E_max_*), demonstrating that these intracellular S/T residues contribute to efficient receptor-mediated G protein activation. Particularly, T312A displayed a modest leftward shift in the concentration-response curve, indicative of an increased apparent agonist potency despite its reduced maximal signaling (Fig. 2C). Additionally, beyond agonist-evoked responses, we also examined basal cAMP production in the absence of agonist. Strikingly, only the helix 8 mutants, S306A and T312A, displayed significantly elevated basal luminescence compared with MC4R^WT^, whereas mutations within the C-terminal tail (S329A, S330A, and the S329A:S330A double mutant) had little effect on constitutive activity (Fig. 2E). Importantly, because all constructs exhibited comparable surface expression, the increase in basal signaling cannot be explained by differences in receptor abundance at the plasma membrane.

**Figure 2:**
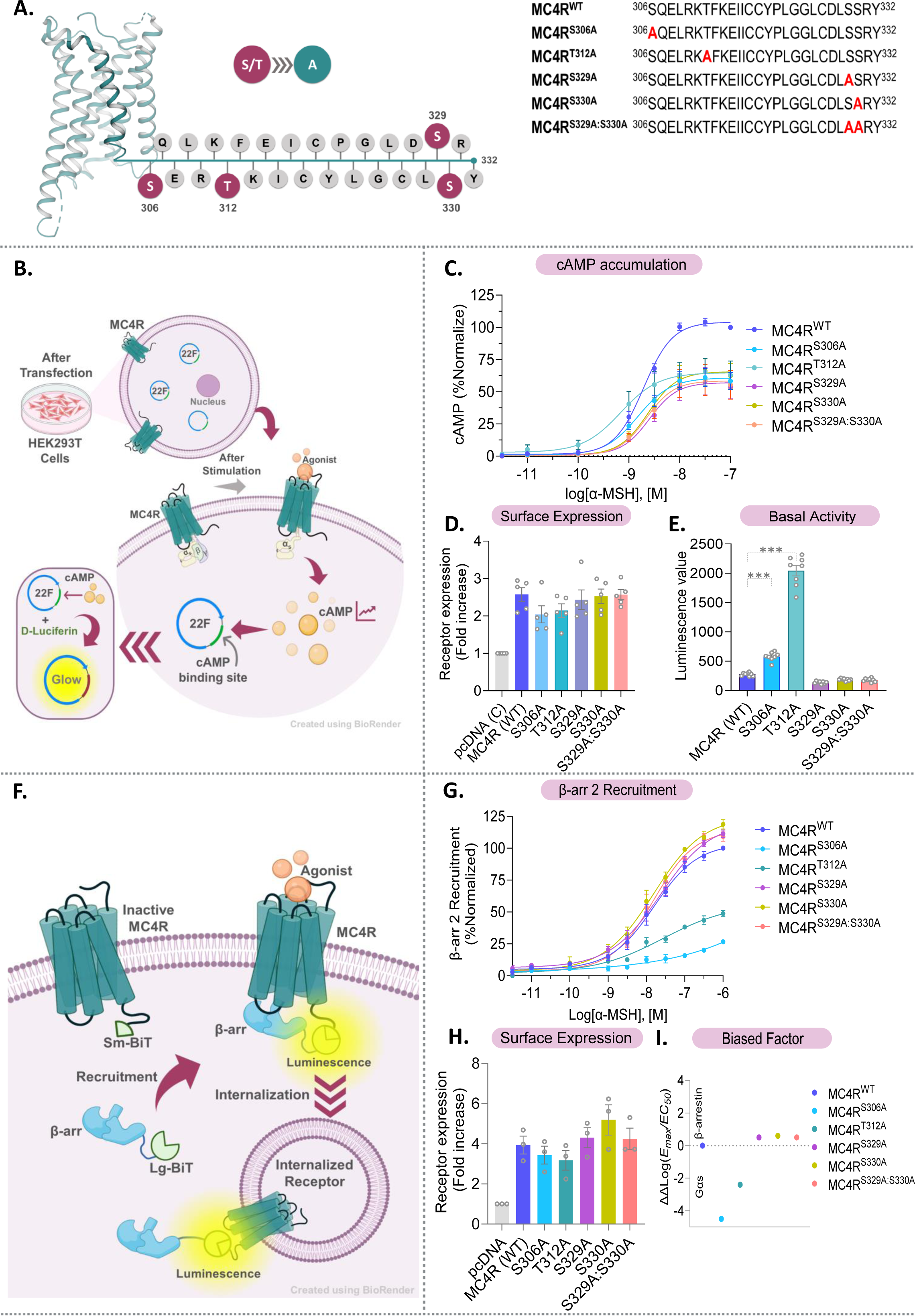
Comparative study with MC4R endogenous ligand (α-MSH) and the respective mutants: **A.** Schematic representation of helix8 and c-terminal of MC4R receptor with phosphorylation sites of S/T residues; along with sequences of the site mutants generated in this study; mutated S/T to Alanine are highlighted in red**; B.** Schematic representation of Gαs activation profiling via luminescence-based cAMP accumulation assay**; C.** α-MSH induced, % normalized of luminescence data for cAMP accumulation in HEK293T cells, 20min, n=5**; D.** Surface expression of MC4R^WT^ & its mutants, for Gα_s_ activation study (n=5), was analyzed using one-way ANOVA with Dunnett test where no significant differences seen in expression**; E.** Basal signaling of MC4R^WT^ & its mutants, n=8, was analyzed using one-way ANOVA with Dunnett test, (***p < 0.001), at 95% CI; **F.** Schematic representation of NanoBiT-based β-arrestin recruitment assay**; G.** α-MSH induced, β-arrestin 2 recruitment in MC4R^WT^ and its mutants, % normalized NanoBiT based luminescence data in HEK293T cells, 10min, n=3**; H.** Surface expression of MC4R^WT^ & its mutants, for β-arr recruitment study part (n=3), was analyzed using one-way ANOVA with Dunnett test where no significant differences seen in expression**; I.** Receptor mediated bias was studied; where, (-ve) value indicates G-protein Biased; 0 indicates balanced while (+ve) value represents β-arrestin biased and the calculated bias factor was plotted.

Together, these data reveal a clear functional distinction between residues located in helix 8 and those within the C-terminal tail. Whereas mutations in the C-terminal phosphosites primarily impair agonist-evoked signaling, substitutions at S306 and T312 additionally increased basal Gα_s_ signaling while reducing the maximal response to α-MSH, consistent with a perturbation of the receptor’s basal conformational equilibrium that favors Gα_s_-competent states but limits the receptor’s capacity to achieve the fully agonist-stabilized signaling state. In contrast, mutations of the C-terminal residues S329 and S330 reduced agonist-stimulated *E_max_* without increasing basal activity, indicating that these residues influence agonist-evoked signaling without producing a detectable shift toward constitutive Gα_s_ activation. Thus, although both H8 and C-terminal mutations impair agonist-induced signaling, their distinct effects on basal activity suggest that these intracellular regions regulate MC4R signaling through mechanistically separable processes.

### 2.3 Helix 8 residues are critical determinants of β-arrestin2 recruitment to MC4R

Having established that residues within both H8 and the C-terminal tail can individually influence Gα_s_-mediated signaling, we next asked whether these regulatory regions similarly control recruitment of βarr, the second major effector arm of MC4R signaling. We quantified agonist-dependent βarr recruitment using a NanoBiT complementation assay, which reports ligand-induced proximity between MC4R and βarr in living cells and integrates receptor-arrestin recruitment across the cellular compartment. So, we first examined the βarr isoform preference of MC4R^WT^. Interestingly, MC4R displayed a pronounced preference for βarr2 over βarr1, with robust α-MSH-dependent βarr2 recruitment but little detectable recruitment of βarr1 under our assay conditions (Fig. S2). This observation is consistent with previous work demonstrating that βarr2 plays a predominant role in agonist-dependent MC4R internalization, with KD of βarr2 producing a substantially greater impairment in MC4R endocytosis than depletion of βarr1(*28*). To further establish that this isoform preference was not an intrinsic limitation of the assay, we performed parallel measurements with GLP-1R, which is well known to engage both βarr isoforms, which readily recruited both βarr1 and βarr2 in our assay (Fig. S2). Together, these experiments established βarr2 as the predominant arrestin isoform engaged by MC4R under our experimental conditions.

We next asked whether the H8 and C-terminal residues identified above, contribute similarly to βarr2 recruitment. Strikingly, the mutations segregated into two distinct and opposing functional phenotypes based on their location within the receptor (Fig. 2G). Substitution of either S306 or T312 within H8 produced a pronounced loss of βarr2 recruitment, with substantial reductions in both apparent potency and maximal recruitment relative to MC4R^WT^. In contrast, mutation of the previously reported C-terminal phosphosites S329 or S330, either individually or in combination, did not impair βarr2 recruitment. Instead, these mutants retained robust agonist-dependent recruitment and displayed a modest increase in maximal response compared with MC4R^WT^ (Fig. 2G). Importantly, these differences could not be readily attributed to altered receptor abundance at the plasma membrane as our whole-cell ELISA performed in parallel demonstrated comparable cell-surface expression across the receptor variants (Fig. 2H). This indicates that the severe impairment in βarr2 recruitment observed for S306A and T312A reflects a functional consequence of disrupting these H8 residues rather than a defect in receptor delivery to the plasma membrane.

Together, these findings reveal an unexpected distinction between the two intracellular regulatory regions of MC4R. Whereas mutation of the canonical C-terminal phosphorylation sites does not compromise βarr2 recruitment, disruption of either H8 residue markedly impairs arrestin engagement and shifts receptor signaling toward a Gs-favored profile (Fig. 2I; Table. S1). Thus, H8, rather than the distal C-terminal tail, emerges as a critical determinant of MC4R-βarr2 coupling suggesting that MC4R may rely more heavily on receptor-core engagement than on its sparse C-terminal phosphorylation sites for arrestin recognition.

### 2.4. Helix 8 disruption impairs MC4R signaling, receptor trafficking, and sustained ERK activation

Having established that disruption of the H8 residues S306 and T312 markedly impairs βarr2 recruitment, we next asked whether the pronounced defect in βarr2 recruitment observed for the H8 mutants translated into altered receptor trafficking. To directly monitor agonist-induced receptor trafficking, we used a NanoBiT-based FYVE assay in which receptor proximity to the FYVE domain reports accumulation of receptor in early endosomal compartments (Fig.4D) Following stimulation with α-MSH, MC4R^WT^ showed a rapid increase in endosomal accumulation, reaching a maximum within approximately 10-20 min before gradually declining. In striking contrast, both H8 mutants exhibited substantially reduced endosomal accumulation. S306A showed the most severe trafficking defect, with T312A displaying an intermediate phenotype (Fig. 4E). The C-terminal mutants, in contrast to H8 mutants, retained considerably greater endosomal trafficking (Fig. 4E). These findings further indicate that H8 mutants particularly the S306, compromises βarr2-dependent receptor trafficking, whereas the C-terminal mutations have a comparatively limited effect on this process. Given that the H8 mutations also markedly altered basal and agonist-induced Gα_s_ signaling, we next asked whether these changes in receptor signaling and trafficking translate into altered downstream MAPK activation. MC4R predominantly signals through Gα_s_, and Gα_s_-dependent signaling has been implicated in MC4R-mediated ERK1/2 activation. Therefore, we first examined agonist-induced ERK1/2 phosphorylation as an early downstream signaling readout (Fig.3A). HEK293T cells expressing MC4R^WT^ or the indicated mutants were stimulated with 1 μM α-MSH for 5 min, followed by immunoblotting for phosphorylated ERK1/2. MC4R^WT^ exhibited a robust increase in ERK1/2 phosphorylation following α-MSH stimulation. In contrast, the H8 mutants displayed markedly attenuated ERK activation, with S306A showing the most pronounced reduction, whereas T312A retained a measurable but reduced response. Notably, T312A showed the greatest preservation of early ERK activation among the H8 mutants, consistent with its relatively stronger G-protein signaling observed in the cAMP assay (Fig. 3, C and D). These findings indicate that disruption of H8 compromises an early downstream MAPK response, extending the functional consequences of these mutations beyond βarr2 recruitment and receptor trafficking.

**Figure 3:**
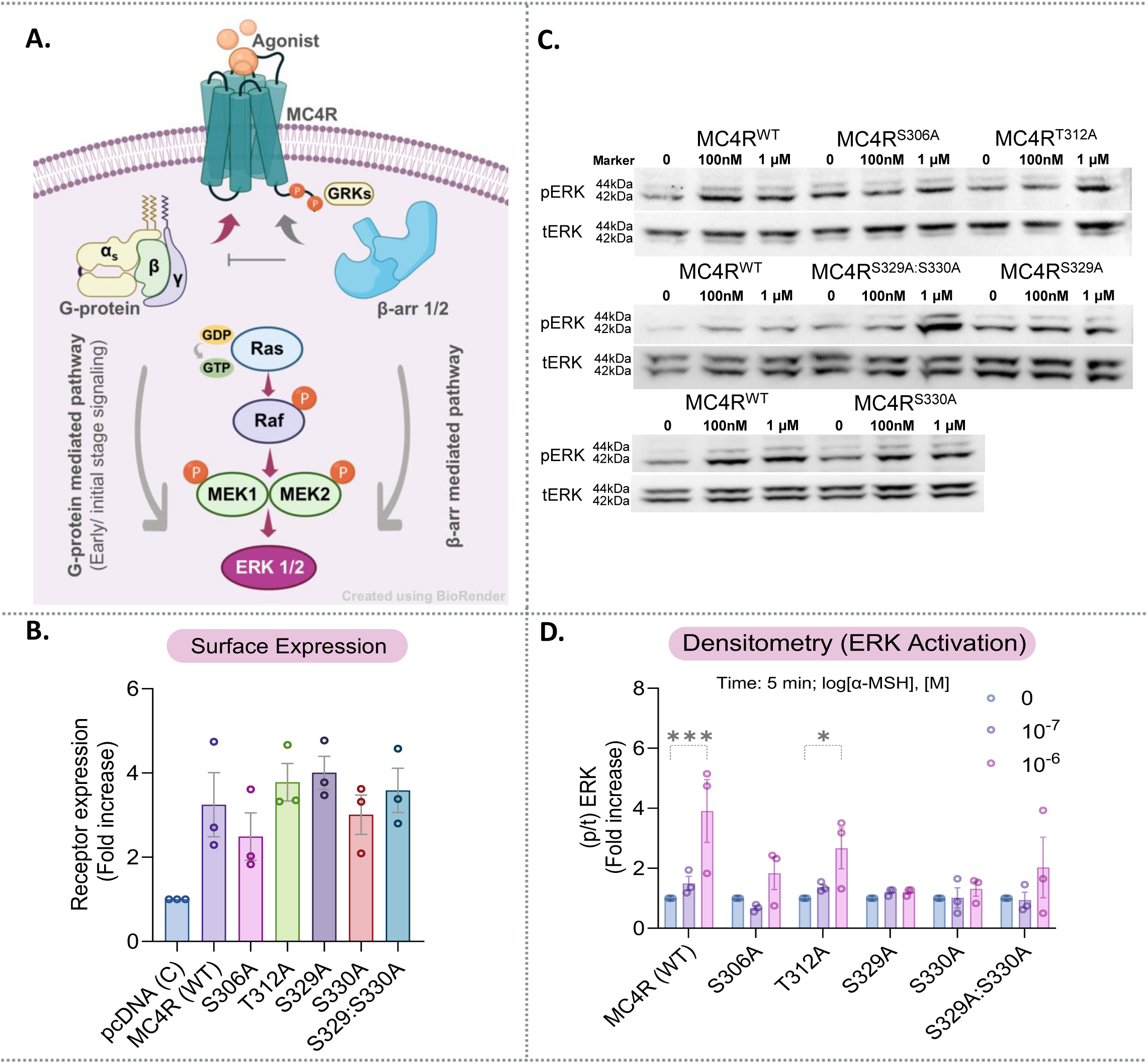
α-MSH-induced phosphorylation of ERK1/2 in HEK-293 cells expressing wild type MC4R and its mutants using Western blot method: **A.** Schematic representation of ERK1/2 activation through G-protein (Early stage) and β-arr mediated.**; B.** Surface Expression, n=3**; C, D.** Western Blot images developed in BioRad’s ChemiDoc system was proceeded with densitometry-based quantification of data (mean ± SEM) from n=3; is presented as bar graphs, normalized with respect to minimal dose under control condition (treated as 1 fold), and analyzed using two-way ANOVA with Dunnett test (***p< 0.001, *p < 0.033); here the error bar indicates ±SEM.

**Figure 4:**
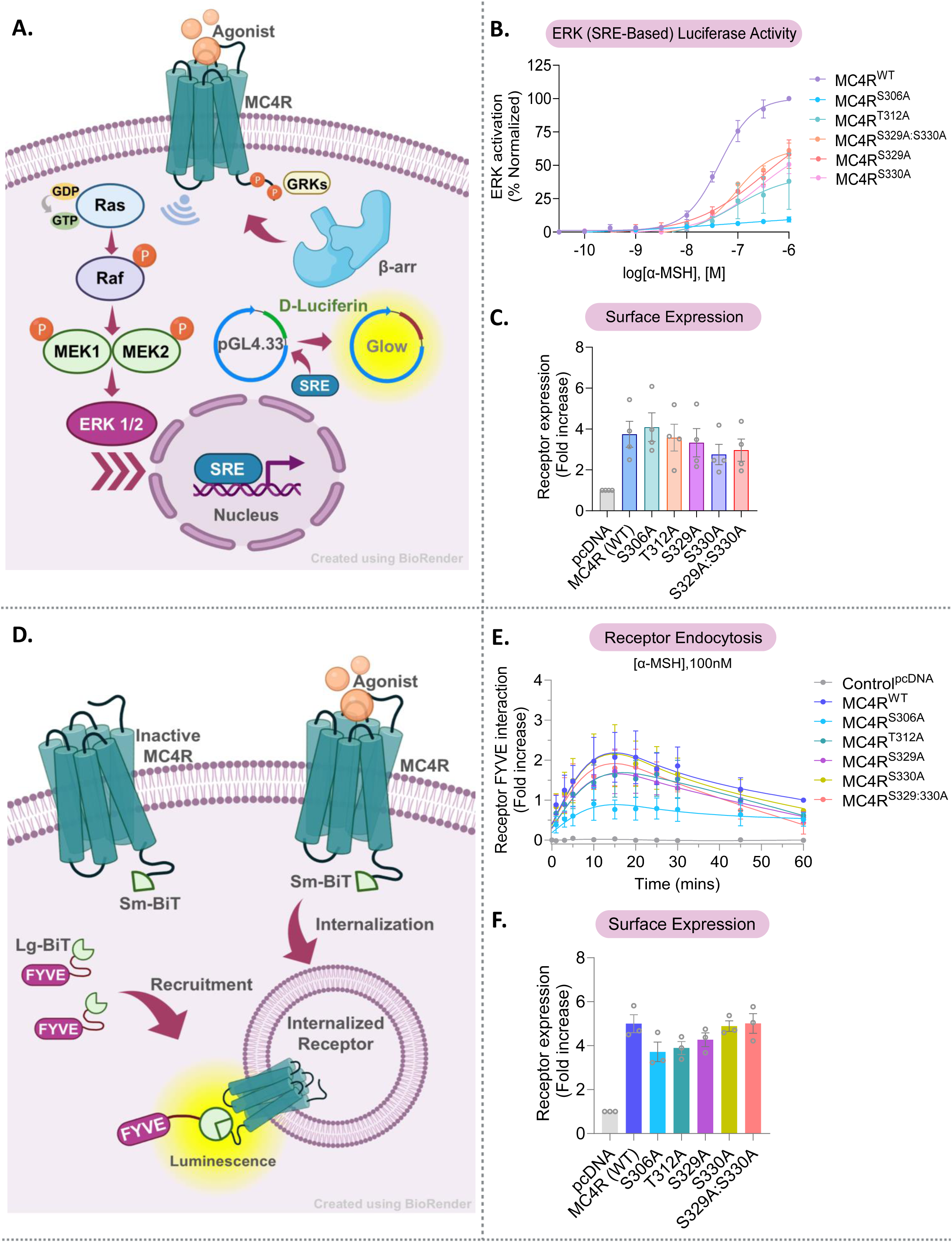
Comparative study with MC4R endogenous ligand (α-MSH) and the respective mutants: **A.** Schematic representation of ERK activation via SRE Reporter luminescence-based assay**; B.** Comparison with MC4R endogenous ligand (α-MSH) and the respective mutants; n=4; % normalized of luminescence data for SRE-ERK activation in HEK293T cells**; C.** Surface expression, n=4, is presented as bar graphs, normalized with respect to control pcDNA under control condition (treated as 1-fold), and analyzed using One-way ANOVA with Dunnett test where no significant differences seen in expression**; D.** Schematic representation of NanoBiT based Receptor Endocytosis assay**; E.** Comparison of endocytosis pattern with 100nM, α-MSH induced, FYVE and MC4R^WT^ and its variant interactions, % normalized NanoBiT based luminescence data in HEK293T cells, n=3**; F.** Surface expression in bar graph, n=3, was analyzed using one-way ANOVA with Dunnett test where no significant differences seen in expression; in every experiment error bar indicates ±SEM.

Since ERK1/2 phosphorylation represents an acute signaling output, we next asked whether the altered receptor signaling produced measurable consequences at a longer timescale. We therefore examined SRE-luciferase activity following prolonged α-MSH stimulation as a transcriptional readout of MC4R signaling (Fig.4A). MC4R^WT^ produced a robust SRE response, whereas the mutants displayed strikingly different transcriptional profiles. The C-terminal mutants retained substantial SRE activity, with the S329A:S330A double mutants showing the highest response among the mutants. In contrast, both H8 mutants were strongly impaired, with S306A exhibiting the weakest transcriptional response (Fig. 4B). Thus, H8 disruption produces a broad functional phenotype spanning proximal receptor activation, βarr2 engagement, receptor trafficking, and longer-timescale signaling. The marked reduction in SRE activity in S306A and T312A mutants, together with their impaired G-protein signaling and βarr2 recruitment, suggests that H8 integrity is required for efficient propagation of MC4R signals from receptor activation to downstream cellular responses. Collectively, these findings establish helix 8 as a critical regulatory element of MC4R signaling and trafficking.

### 2.5. Helix 8 and C-terminal residues differentially regulate MC4R transducer coupling

The effects of H8 mutants therefore extend beyond individual signaling readouts and are evident across multiple stages of the MC4R response, from proximal receptor activation and βarr2 recruitment to receptor trafficking and sustained ERK-dependent transcription. These findings prompted us to further probe if the consequences of H8 disruption restricted to the canonical Gαs/βarr2 signaling axis, or whether individual residues also influence the receptor’s ability to engage alternative intracellular transducers. Although MC4R signaling is predominantly mediated through Gα_s_, the receptor is capable of engaging multiple G-protein families, including Gα_q/11_ and Gα_12/13_, thereby expanding the functional repertoire of MC4R signaling (*29*, *30*). Having observed that disruption of H8 markedly altered Gα_s_ signaling and βarr2 recruitment, we next asked whether these effects were restricted to these canonical signaling pathways or instead reflected a broader alteration in the receptor’s ability to engage distinct intracellular transducers. We therefore examined agonist-induced activation of NFAT-RE and SRF-RE reporters as transcriptional readouts associated with Gα_q/11_-and Gα_12/13_-dependent signaling, respectively. The mutations produced distinct effects across these pathways. In the NFAT-RE assay, MC4R^WT^ produced the strongest response, whereas S329A retained a substantial fraction of receptor activity. In contrast, S306A, T312A, S330A, and the S329A:S330A double mutant showed markedly reduced NFAT responses, with S330A exhibiting the strongest impairment (Fig. 5C). Thus, individual regulatory residues differentially influence MC4R signaling through the Gα_q/11_-associated pathway. A distinct pattern was observed for SRF-RE signaling. MC4R^WT^ again produced the strongest response, whereas T312A retained substantial activity despite its pronounced defects in βarr2 recruitment and sustained ERK signaling. In contrast, S306A, S329A, S330A, and the S329A:S330A double mutant displayed strongly attenuated SRF-RE responses (Fig. 5D). Notably, S330A showed a particularly strong reduction in both NFAT-RE and SRF-RE responses while retaining substantial SRE activity, suggesting that this C-terminal residue contributes differentially to signaling through distinct transducer pathways. Conversely, the relative preservation of Gα_12/13_-associated signaling in T312A, compared to βarr2 recruitment and sustained ERK signaling, further highlights the pathway-selective consequences of perturbing individual regulatory residues. Together, these findings indicate that the MC4R H8 and C-terminal regions do not simply determine overall receptor activity but contribute to the differential engagement of distinct intracellular signaling pathways. Thus, individual residues within these regulatory regions may fine-tune the receptor’s signaling output by differentially influencing transducer-specific coupling and downstream pathway propagation.

**Figure 5:**
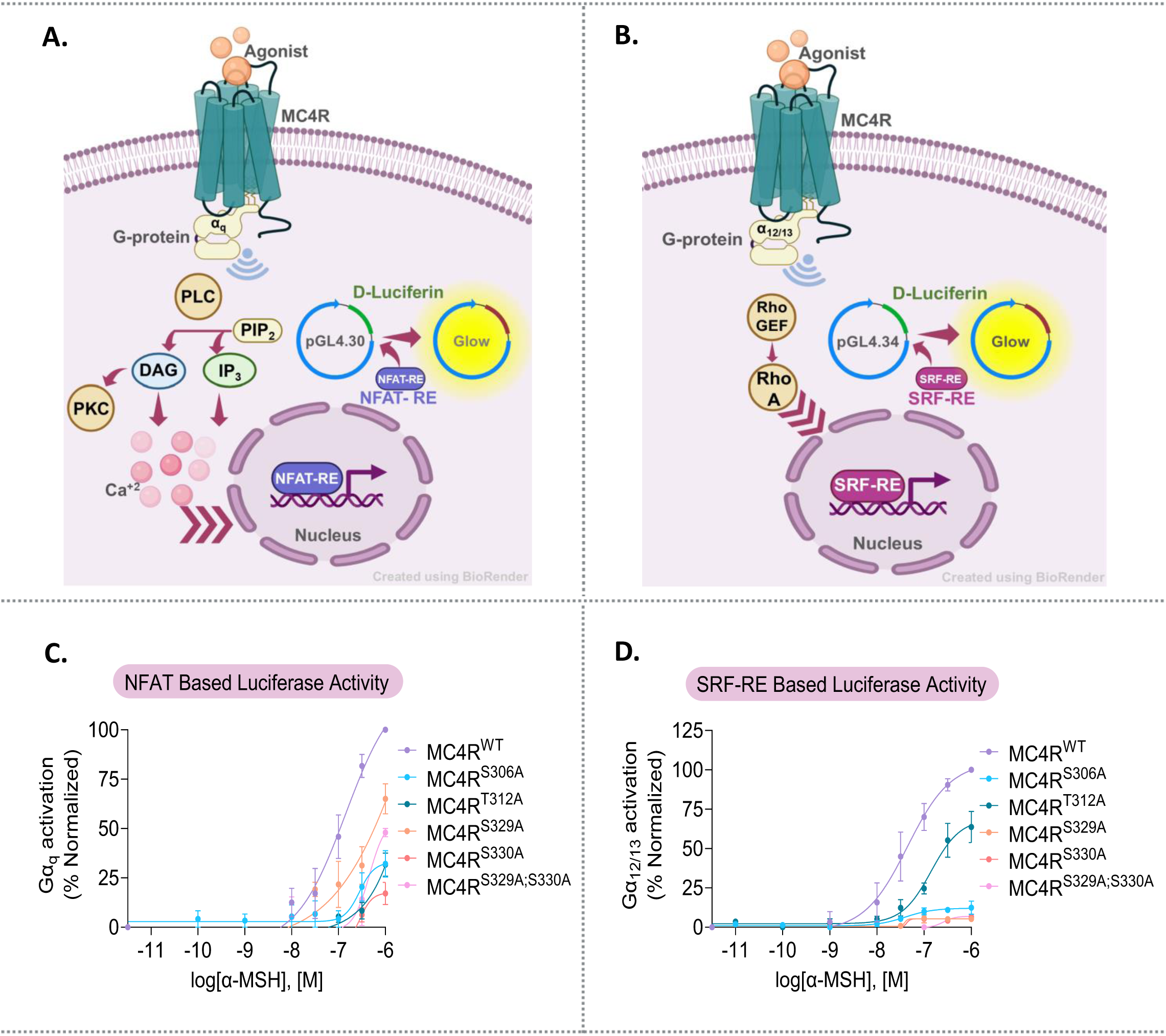
Comparative study with MC4R endogenous ligand (α-MSH) and the respective mutants: **A.** Schematic representation of Gα_q_ activation via NFAT-RE Reporter luminescence-based assay**; B.** Schematic representation of Gα_12/13_ activation via SRF-RE Reporter luminescence-based assay; **C.** Gα_q_ activation via NFAT-RE was observed, α-MSH induced, n=3; % normalized of luminescence data in HEK293T cells; **D.** Gα_12/13_ activation via SRF-RE was observed, α-MSH induced; % normalized of luminescence data in HEK293T cells, n=3; in both experiment error bar indicates ±SEM.

### 2.6 Ligand-Specific Pharmacological Remodeling of MC4R Regulatory Site Mutants

Having established that the H8 and C-tail mutants differentially affect transducer specific transcriptional response downstream of the MC4R, we next asked whether these defects represent an intrinsic inability of the mutant receptors to adopt productive signaling conformations or could be differentially modulated by the agonist bound to the receptor. This distinction is particularly important for MC4R, where chemically diverse agonists can engage distinct receptor conformations and produce divergent signaling outcomes. We therefore expanded our analysis beyond the endogenous agonist α-MSH to a pharmacologically diverse panel comprising the peptide agonists Melanotan-I (MT-I), Melanotan-II (MT-II), and bremelanotide, together with the structurally distinct small-molecule agonist THIQ. Rather than simply assessing whether these ligands retained activity at the mutant receptors, we compared their potency and maximal efficacy across Gα_s_-mediated cAMP accumulation and β-arrestin2 recruitment. This allowed us to determine whether the signaling defects imposed by these regulatory site mutants were ligand-independent, reflecting a generalized impairment in receptor activation, or ligand-dependent, revealing agonist-specific opportunities to stabilize productive signaling states.

#### 2.6.1 Ligand-dependent remodeling of Gα_s_ signaling

We first examined Gα_s_-mediated cAMP accumulation across the receptor variants. At MC4R^WT^, the synthetic agonists displayed greater functional potency than α-MSH, with bremelanotide exhibiting the highest potency, followed by MT-II, MT-I, THIQ, and α-MSH. This overall increase in functional potency of the synthetic agonists was largely retained across the H-8 and C-terminal mutants, indicating that these substitutions did not uniformly abolish the ability of the receptor to respond to chemically diverse agonists. In contrast to the relatively conserved potency profiles, maximal cAMP responses exhibited a pronounced ligand-and mutation-dependent pattern. Whereas THIQ produced the highest efficacy at MC4R^WT^ and S329A, MT-II produced the greatest response at T312A and S329A:S330A, while MT-I produced the highest response at S330A (Fig. 6, A to H). Thus, the mutations did not simply reduce the overall signaling capacity of MC4R, rather, they altered the relative efficacy with which individual agonists engaged the Gα_s_ pathway. These findings reveal ligand-dependent remodeling of the receptor-efficacy relationship, suggesting that chemically distinct agonists can differentially stabilize signaling-competent states of MC4R carrying distinct regulatory-region mutation.

**Figure 6:**
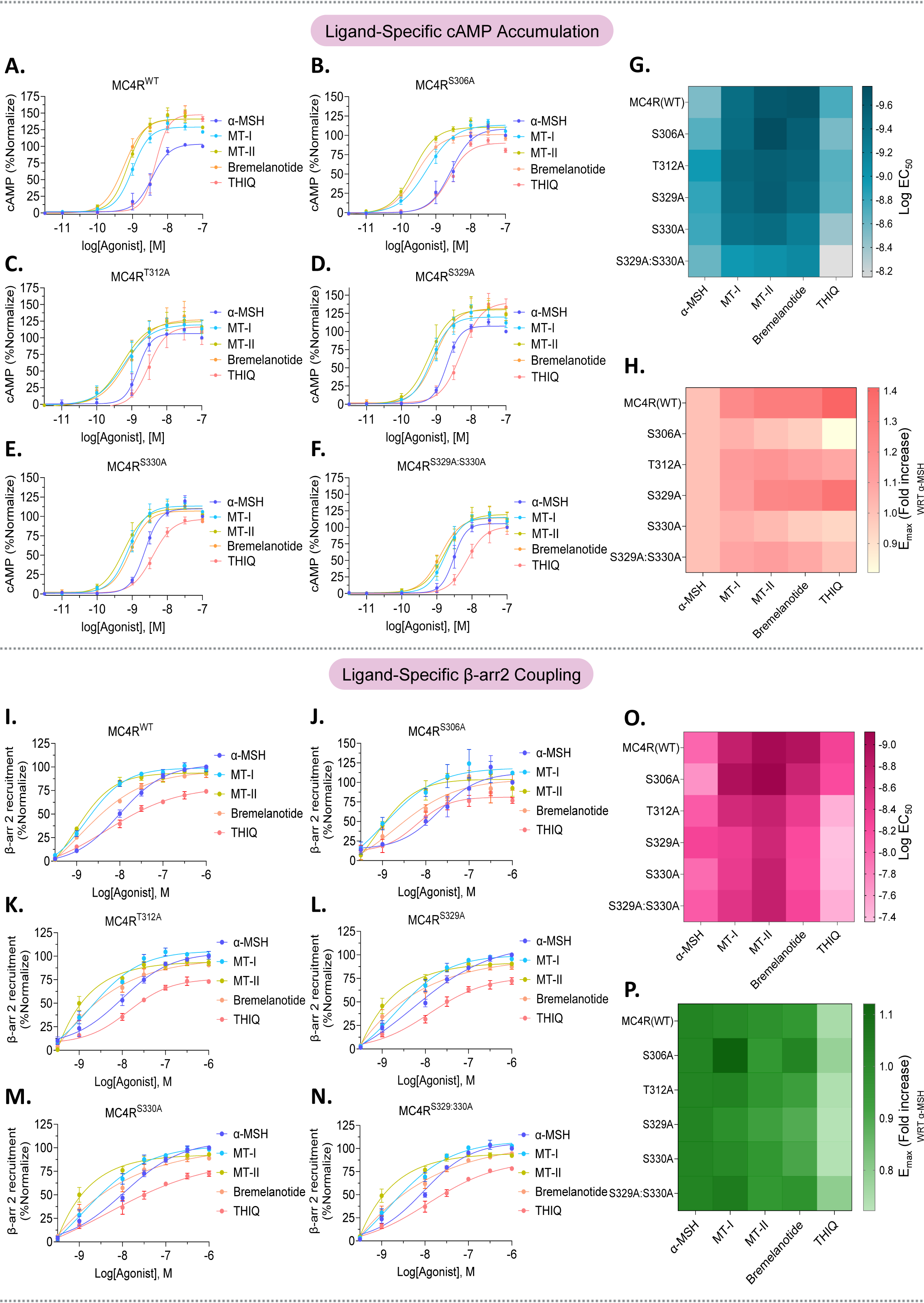
Ligand-Specific Pharmacological Profiling of MC4R Variants; Gα_s_ Activation and cAMP Accumulation Profile comparison with **A.** MC4R^WT^**; B.** MC4R^S306A^**; C.** MC4R^T312A^**; D.** MC4R^S329A^**; E.** MC4R^S330A^**; F.** MC4R^S329A:S330A^; and the respective pharmacological agents; n=3; % normalized luminescence data for cAMP accumulation, in HEK293T cells**; G, H.** Comparative analysis was made and presented in the form of Heat MAP for LogEC_50_ and E_max_ values obtained from Gα_s_ activation assay; Modulation of β-Arrestin 2 Recruitment comparison with **I.** MC4R^WT^**; J.** MC4R^S306A^**; K.** MC4R^T312A^**; L.** MC4R^S329A^**; M.** MC4R^S330A^**; N.** MC4R^S329A:S330A^; and the respective pharmacological agents; n=4; % normalized NanoBiT based luminescence data for β-arrestin 2 recruitment, in HEK293T cells**; O, P.** Comparative analysis was made and presented in the form of Heat MAP for LogEC_50_ and E_max_ values obtained from β-arrestin 2 recruitment assay; in every experiment error bar indicates ±SEM.

#### 2.6.2 Ligand-dependent modulation of β-arrestin2 recruitment

We next asked whether the ligand-dependent remodeling observed for Gα_s_ signaling was also evident in βarr2 recruitment. At MC4R^WT^, MT-II exhibited the highest potency for β-arr2 recruitment, followed by Bremelanotide, MT-I, and α-MSH, whereas THIQ displayed substantially lower maximal efficacy, consistent with partial agonism in this pathway (Fig. 6I to P). The H8 and C-tail mutants retained the ability to recruit βarr2 in response to the synthetic agonists, but the magnitude of recruitment was differentially affected by both the mutation and the ligand. This was particularly evident for S306A. Although α-MSH produced a marked reduction in βarr2 recruitment at S306A, MT-I elicited substantially greater recruitment and produced the highest maximal response among the agonists tested at this mutant (Fig. 6J). Thus, the βarr2 defect associated with S306A was not uniformly refractory to agonist stimulation but could be partially alleviated by an alternative ligand. Moreover, the ability of MT-I to produce greater βarr2 recruitment at S306A than α-MSH suggests that agonist structure can influence the extent to which the altered receptor engages a βarr2-competent signaling state. Therefore, the functional consequences of the S306A substitution are not solely determined by the receptor sequence itself but emerge from an interaction between receptor state and ligand identity.

Although these analyses revealed ligand-and mutation-dependent differences in individual signaling outputs, comparison of pathway-specific *E_max_* and potency alone does not establish how these changes reshape the relative coupling preference of the receptor. We therefore integrated the Gα_s_ and β-arrestin2 pharmacological profiles using an operational model of ligand-directed bias to determine whether the mutations altered the signaling preference of MC4R and whether this phenotype could be modulated by agonist structure.

### 2.7 Ligand-Directed Bias Reveals Pharmacological Remodeling of Mutant Phenotypes

Having established ligand-and mutation-dependent differences in Gα_s_ and βarr2 signaling, we next integrated these pathway-specific responses using a relative bias analysis based on ΔΔlog(*E_max_*/*EC_50_*) (Table. S3 to S8), with α-MSH serving as the reference ligand for each receptor variant. At MC4R^WT^, the synthetic agonists produced only modest deviations from the α-MSH reference, with a slight overall tendency toward G-protein-favored signaling (Fig. 7, A to F). Strikingly, the direction and magnitude of ligand-dependent bias were markedly altered by the H8 and C-tail mutants. At S306A, all synthetic agonists shifted the signaling profile toward βarr relative to α-MSH, with THIQ producing the largest positive shift. This relatively-βarr-directed shift is consistent with the enhanced β-arr2 recruitment observed for selected synthetic agonists at S306A despite the impaired response to α-MSH. In contrast, S329A exhibited the opposite pattern. All synthetic agonists produced substantial negative ΔΔlog(*E_max_*/*EC_50_*) values, indicating a pronounced shift toward G-protein-favored signaling relative to α-MSH. Thus, the same pharmacological panel produced opposing changes in relative signaling preference depending on the regulatory-site mutations. The remaining variants showed comparatively modest shifts. T312A and S330A remained close to the reference state, with only mild G-protein-directed effects for selected agonists, whereas the S329A:S330A double mutant displayed predominantly modest deviations from α-MSH. Collectively, these data demonstrate that ligand-directed signaling bias is strongly dependent on receptor sequence, rather than imposing a fixed signaling preference, different regulatory-site substitutions reshape how individual agonists redistribute signaling between Gα_s_ and β-arrestin2. The bias analysis therefore revealed that phosphorylation-site mutations do not impose a uniform signaling preference. Instead, the relative contribution of Gα_s_ and βarr2 signaling could be substantially reshaped by agonist structure in a mutation-dependent manner. However, these analyses were based on proximal receptor-effector responses. We next asked whether the ligand-and mutation-dependent remodeling observed at this level was propagated to a downstream functional signaling output.

**Figure 7:**
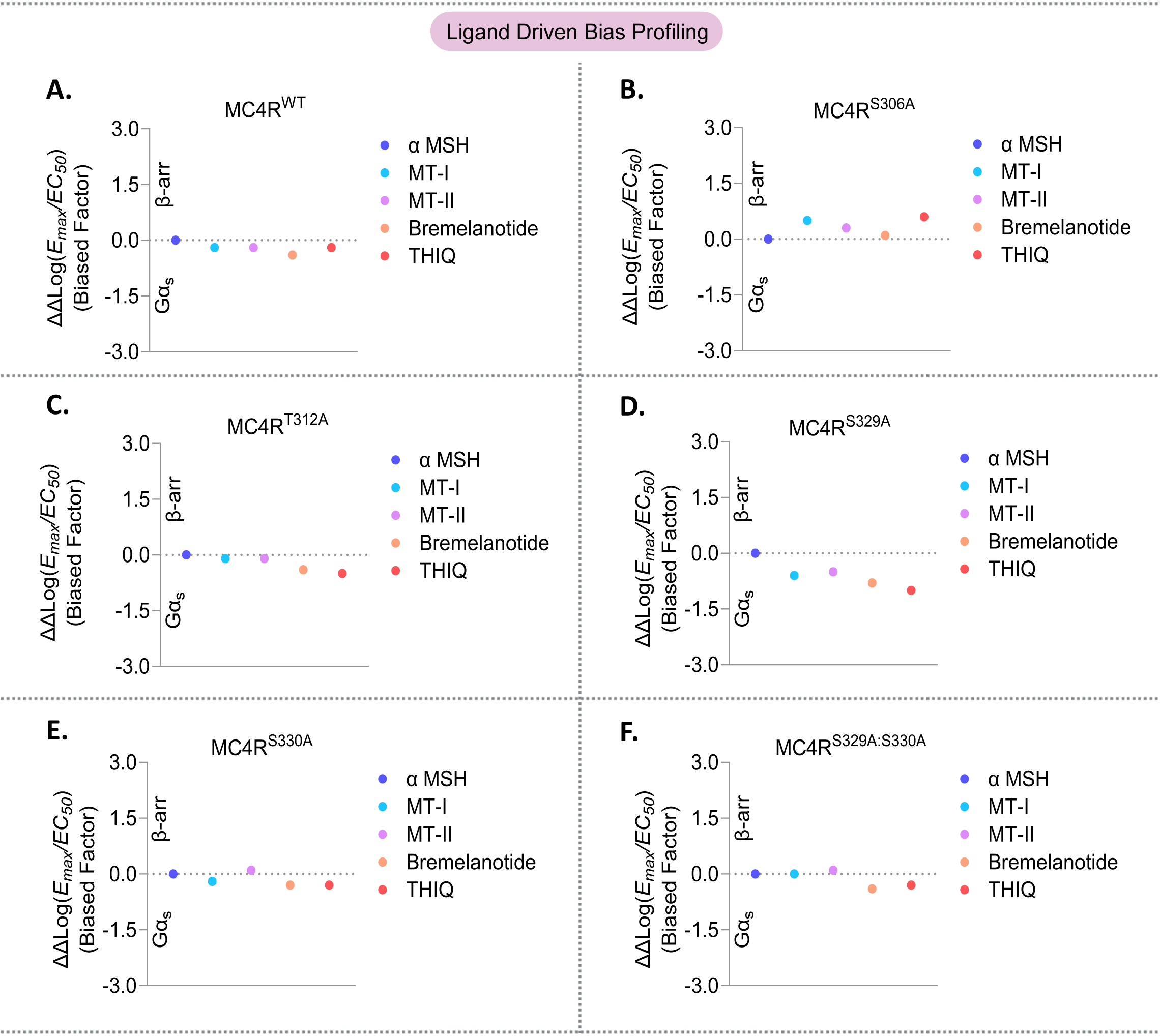
B**i**ased **Factor determination of ligands with all the variants:** Scatter plot comparing pharmacological agents w.r.t. its endogenous ligand α-MSH; here (-ve) value indicates G-protein Biased; 0 indicates balanced while (+ve) value represents β-arrestin biased. So Comparative analysis was made with **A.** MC4R^WT^**; B.** MC4R^S306A^**; C.** MC4R^T312A^**; D.** MC4R^S329A^**; E.** MC4R^S330A^**; F.** MC4R^S329A:S330A^; and the respective pharmacological agents by calculating the bias factor.

### 2.8 Propagation of Ligand-Dependent Signaling Remodeling to Downstream MAPK/Transcriptional Signaling

To address this question, we examined serum response element (SRE)-dependent transcriptional activity, which provides an integrated readout of sustained MAPK signaling downstream of MC4R activation. This allowed us to determine whether the altered pharmacological hierarchy observed at the receptor-proximal level was preserved or further remodeled at a downstream signaling node (Fig. 8, A to H). At MC4R^WT^, the synthetic agonists displayed greater potency than α-MSH, with MT-II > MT-I > bremelanotide > THIQ. However, their efficacy hierarchy differed substantially, with THIQ and bremelanotide producing lower maximal transcriptional responses than the peptide agonists. This distinction between potency and efficacy became markedly more pronounced in the regulatory-site mutants. Most strikingly, S306A exhibited a pronounced reordering of ligand efficacy. Whereas THIQ produced the weakest maximal transcriptional response at MC4R^WT^, it elicited the highest response at S306A. Conversely, the potency hierarchy was also remodeled, with MT-I producing the most potent response and MT-II the least potent response at this mutant. Thus, the S306A mutation did not simply attenuate downstream signaling; rather, it changed the relative functional efficacy with which different agonists engaged the downstream signaling cascade. Ligand rank ordering was similarly remodeled, although to different extents, across the remaining variants. MT-I was the most potent agonist for T312A, S330A, and S329A:S330A, whereas S329A showed a modestly distinct profile with MT-II displaying the greatest potency. These differences were accompanied by mutation-dependent changes in maximal efficacy, demonstrating that the pharmacological hierarchy established at MC4R^WT^ was not preserved across the various mutants. Together, these findings indicate that receptor mutation and ligand structure jointly determine the functional signaling output generated downstream of MC4R. Importantly, agonist efficacy therefore cannot be considered an intrinsic property of the ligand rather, it emerges from the interaction between ligand chemotype and the signaling landscape of the receptor. Additionally, these findings demonstrate that receptor mutation and ligand structure jointly determine the signaling output propagated downstream, revealing a level of pharmacological plasticity that is not apparent from the wild-type receptor alone. Furthermore, when comparing all receptor variants against each specific ligand, MC4R^WT^ typically elicited the maximal SRE-mediated ERK response (Fig. S3). However, upon stimulation with THIQ, the S329A and S329A:S330A mutants demonstrated slightly greater maximal efficacy than the wild type.

**Figure 8:**
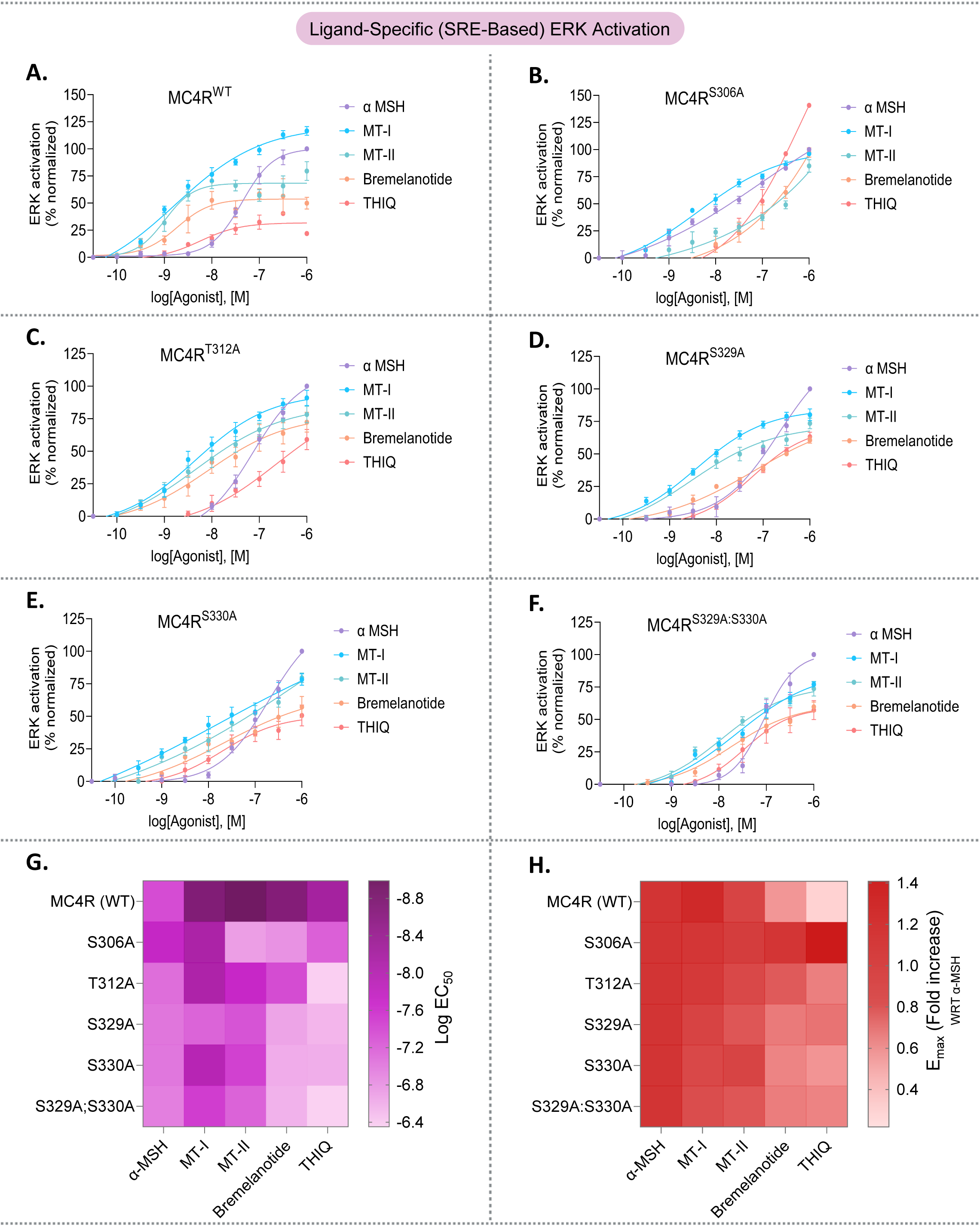
Ligand-Specific ERK Profiling with MC4R Variants: Comparison with MC4R variants, **A.** MC4R^WT^**; B.** MC4R^S306A^**; C.** MC4R^T312A^**; D.** MC4R^S329A^**; E.** MC4R^S330A^**; F.** MC4R^S329A:S330A^; and set of pharmacological agents for ERK activation via luminescence-based SRE reporter assay; n=4; % normalized of luminescence data in HEK293T cells; error bar indicates ±SEM**; G, H.** Comparative analysis was made and presented in the form of Heat MAP for LogEC_50_ and E_max_ values obtained from SRE mediated ERK activation assay.

### 2.9 Pharmacological Constraints of Gα_q/11_ and Gα_12/13_ Dependent Signaling

The ligand-and mutation-dependent remodeling observed for Gα_s_ and βarr2, together with the marked reordering of downstream SRE responses, raised the question of whether this pharmacological plasticity extends uniformly across the MC4R signaling repertoire. We therefore challenged the mutant variants with the same structurally diverse agonist panel to determine whether ligand structure could similarly modulate transcriptional response associated with the Gα_q/11_ and Gα_12/13_ pathways. For Gα_q/11_ signaling (Fig. 9, A to F), MC4R^WT^ exhibited robust responses across the agonist panel, whereas all H8 and C-tail mutants displayed substantially attenuated maximal signaling. Importantly, the mutants were not completely insensitive to ligand identity: individual agonists produced measurable differences in the residual Gα_q/11_ responses, indicating that ligand structure retained some capacity to modulate signaling even in the altered receptors. Nevertheless, none of the synthetic agonists restored mutant responses to the functional levels observed for MC4R^WT^. Thus, in contrast to the pronounced ligand-dependent remodeling observed for Gα_s_ and βarr2, pharmacological modulation of Gα_q/11_ coupling remained constrained largely to the residual signaling capacity of the mutant receptors. Furthermore, ligand specific comparative analysis of Gα_q/11_ signaling (Fig. S4) demonstrated that, unlike the wild-type, the T312A, S330A, and S329A:S330A mutants achieved higher potency and efficacy with bremelanotide than with α-MSH, suggesting that bremelanotide provides an enhanced signaling outcome in these variants. An even more restrictive signaling phenotype was observed for Gα_12/13_ (Fig. 9, G to L). MC4R^WT^ showed robust activation across the agonist panel, whereas most mutants exhibited severely attenuated responses. T312A retained the greatest residual Gα_12/13_ signaling among the mutants, demonstrating that disruption of this residue does not completely abolish coupling to this pathway; however, its maximal response remained substantially below that of MC4R^WT^. The remaining variants showed only weak residual signaling despite stimulation with chemically distinct agonists, indicating that ligand-dependent modulation was insufficient to restore efficient Gα_12/13_ coupling. Additionally, a ligand-specific comparative analysis of Gα_12/13_ signaling (Fig. S5) revealed that all tested pharmacological analogs induced a stronger response than α-MSH in the MC4R-WT variant; while THIQ yielded the highest maximal efficacy across the entire panel of ligands when stimulating the S306A mutant.

**Figure 9:**
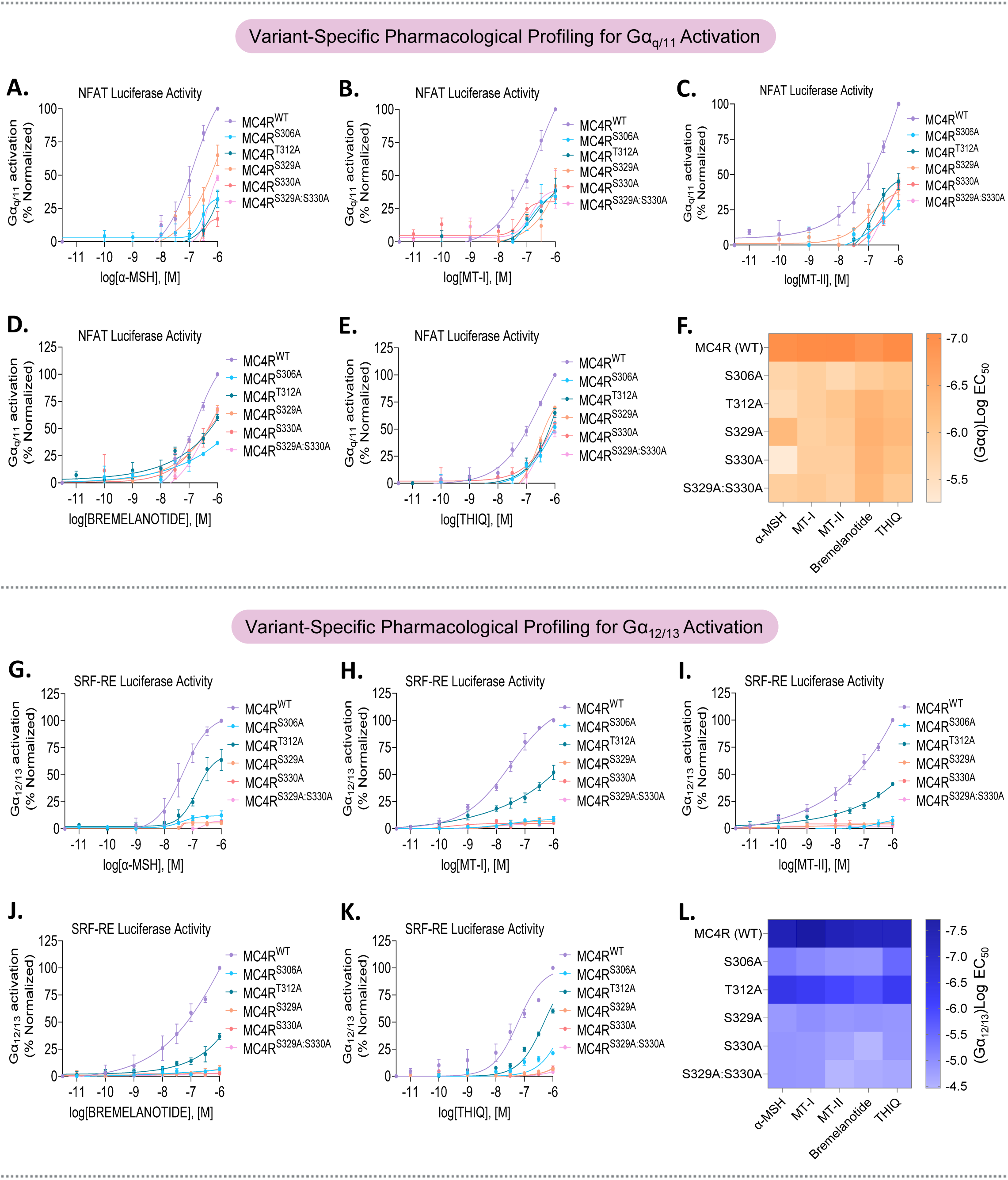
Variant-Specific Gα_q/11_ and Gα_12/13_ Profiling with a panel of Pharmacological Agents; Gα_q_ Profile comparison with **A.** α-MSH**; B.** MT-I**; C.** MT-II**; D.** BREMELANOTIDE**; E.** THIQ; and the respective MC4R variants; n=3; % normalized luminescence data for Gα_q/11_ activation via NFAT-RE reporter assay, in HEK293T cells**; F.** Comparative analysis was made and presented in the form of Heat MAP for LogEC_50_ values obtained from Gα_q/11_ activation; Gα_12/13_ activation comparison with **G.** α-MSH**; H.** MT-I**; I.** MT-II**; J.** BREMELANOTIDE**; K.** THIQ; and the respective pharmacological agents; n=3; % normalized luminescence data for Gα_12/13_ activation via SRF-RE reporter assay, in HEK293T cells**; L.** Comparative analysis was made and presented in the form of Heat MAP for LogEC_50_ values obtained from Gα_12/13_ activation; here in every experiment error bar indicates ±SEM.

Together, these findings reveal that the pharmacological plasticity of MC4R regulatory-site mutants is strongly transducer-and residue-dependent. Whereas ligand structure can substantially remodel Gα_s_ and βarr2 signaling and reorder downstream SRE transcriptional responses, NFAT-RE and SRF-RE responses associated to Gα_q/11_ and Gα_12/13_ remained more strongly constrained following disruption of the H8 and C-terminal residues. Thus, ligand binding can modulate signaling through the altered receptor, but the extent to which this modulation is propagated into downstream transcriptional responses depends on the structural and functional requirements of the individual signaling pathway.

### 2.10 Clinical Agonist Setmelanotide Reveals Pathway-Selective Pharmacological Rescue of MC4R Mutants

Having established that ligand structure can remodel the signaling output of MC4R phosphorylation-site mutants, but that this plasticity is strongly pathway-dependent, we next asked whether this principle extends to a clinically relevant MC4R agonist. The preceding pharmacological profiling revealed two important features of the mutant receptors. First, chemically diverse agonists could substantially reshape signaling through Gα_s_, β-arrestin2, and downstream ERK, demonstrating that the mutations do not simply render the receptor functionally inactive but instead alter the conformational landscape available to different ligands. Second, this pharmacological plasticity was not universal as Gα_q/11_ and Gα_12/13_ associated transcriptional response was profoundly constrained despite stimulation with structurally distinct agonists. We therefore reasoned that an agonist optimized for therapeutic activation of MC4R might exploit this residual conformational plasticity even more effectively. To test this possibility, we extended our analysis to Setmelanotide, an FDA-approved, highly potent MC4R agonist used clinically for chronic weight management (*31*, *32*). Notably, Setmelanotide has previously been reported to preferentially engage the Gα_q/11_ associated signaling pathway at MC4R (*33*), providing a pharmacologically distinct ligand with which to examine whether ligand-specific pathway preference is retained or remodeled across receptor variants. We, therefore, systematically profiled its activity across the same signaling repertoire, encompassing Gα_s_-mediated cAMP accumulation, βarr2 recruitment, downstream ERK signaling, and Gα_q/11_-and Gα_12/13_-dependent NFAT-RE and SRF-RE responses.

Setmelanotide produced its most striking effect on the canonical Gα_s_-cAMP pathway (Fig. 10A). At MC4R^WT^, Setmelanotide produced a pronounced increase in potency, shifting the EC_50_ from approximately 1.9 nM for α-MSH to 0.06 nM. More importantly, the drug substantially remodeled the functional phenotype of the mutant receptors. Unlike α-MSH, which generally revealed reduced signaling capacity across the mutants, Setmelanotide produced enhanced potency at most variants and markedly increased maximal efficacy in several mutants. Notably, S330A represented an exception in which the mutation remained functionally restrictive, whereas the remaining variants displayed greater potency than WT. Furthermore, with the exception of T312A and S330A, the mutant receptors reached higher maximal cAMP responses than WT MC4R. Thus, Setmelanotide does not merely retain activity at the receptor variants, rather, its chemical scaffold can engage receptor conformations that enable substantial, and in some cases enhanced, Gα_s_ signaling despite the underlying structural perturbation (Fig. 10C; Table. S2). Interestingly, the behavior of Setmelanotide in the β-arrestin2 and downstream ERK pathways was markedly different (Fig. 10, B and D). Although Setmelanotide produced a pronounced increase in potency across the receptor variants, it largely preserved the relative efficacy established with α-MSH. In other words, the drug amplified the signaling capacity of the receptors without uniformly erasing the mutation-dependent phenotypes. A similar pattern was observed in the downstream ERK response, where Setmelanotide enhanced signaling potency but did not produce the dramatic functional inversion observed in the Gα_s_ pathway. These observations are important because they demonstrate that the exceptional pharmacological activity of Setmelanotide is not equivalent to global restoration of receptor function. Instead, its effect depends on the signaling axis being interrogated, with the strongest functional compensation occurring within the canonical Gα_s_ pathway.

**Figure 10:**
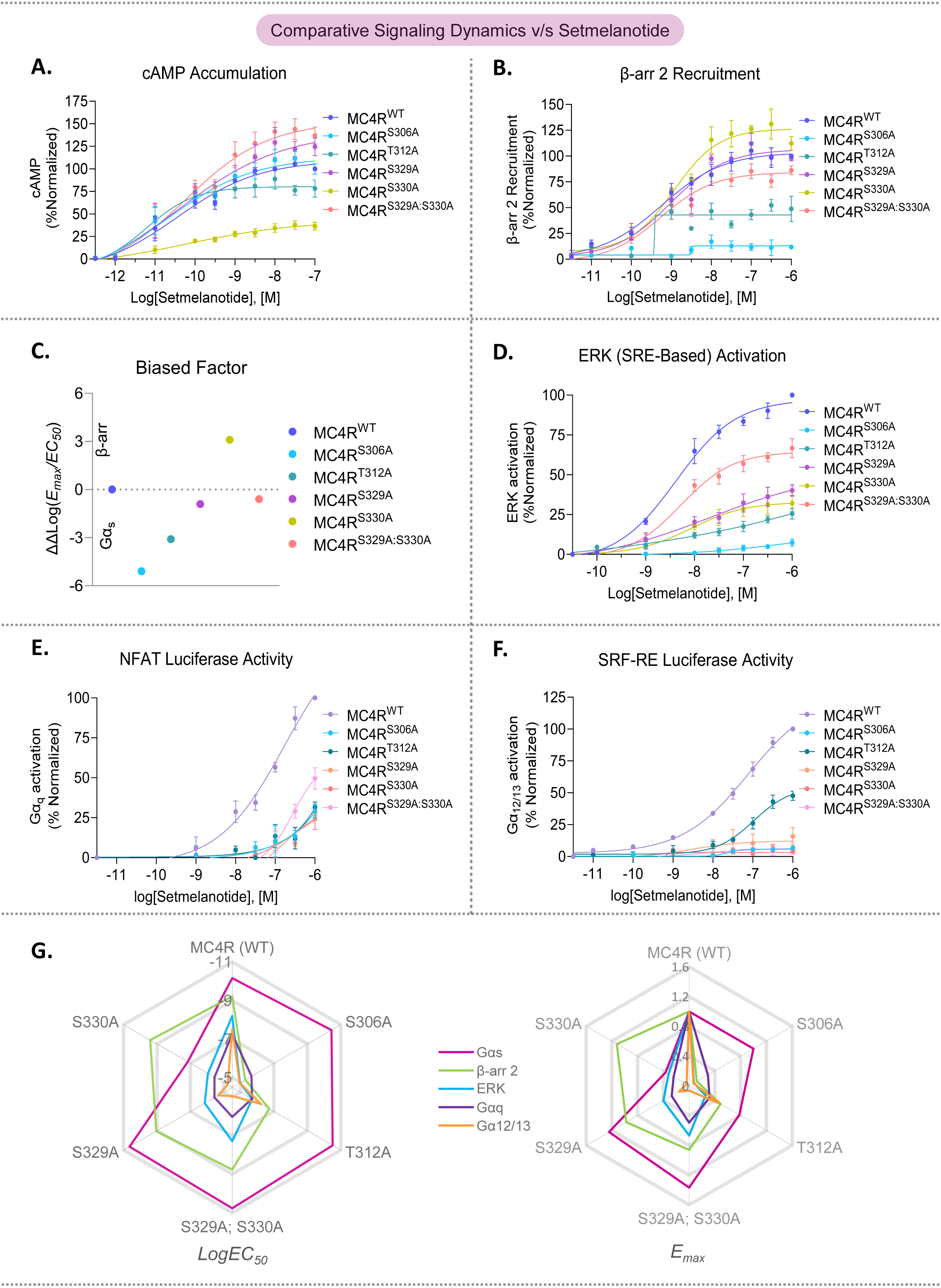
Comparison study with FDA approved ligand (Setmelanotide) and the respective mutants: **A.** Gα_s_ activation profiling via luminescence-based cAMP accumulation assay**; B.** β-arrestin 2 recruitment via NanoBiT based assay **C.** Shows the change in (Gα_s_/β-arr) signaling bias occurred upon Setmelanotide introduction**. D.** ERK activation via SRE Reporter luminescence-based assay**; E.** Gα_q_ activation via NFAT-RE Reporter luminescence-based assay**; F.** Gα_12/13_ activation via SRF-RE Reporter luminescence-based assay; n=3; % normalized of luminescence data in HEK293T cells; in every experiment error bar indicates ±SEM. **G.** Gives a comparative analysis of LogEC_50_ and E_max_ value simultaneously for each signaling profile with each of the variants which is plotted in a radar plot.

This pathway selectivity became even more apparent when we examined the alternative G-protein associated transcriptional responses (Fig. 10, E and F). Despite its exceptional activity through Gα_s_, Setmelanotide did not overcome the marked attenuation of NFAT-RE and SRF-RE responses observed for the receptor variants. The drug produced the expected increase in potency at MC4R^WT^, but none of the mutant receptors showed meaningful restoration of Gα_q/11_ signaling. Likewise, although T312A retained measurable residual Gα_12/13_ activity, Setmelanotide did not restore this response to WT-like levels, while the other mutants remained severely impaired. Thus, even a clinically optimized MC4R agonist cannot bypass the structural requirements imposed by these H8 and C-terminal residues for productive coupling to these alternative transducers.

Taking together, the complete pharmacological analysis reveals that phosphorylation-site mutations do not simply reduce MC4R activity; they reshape the receptor’s pharmacological landscape in a ligand-and transducer-dependent manner. Distinct agonists can differentially exploit the altered conformational ensemble of the mutant receptor, producing substantial remodeling of Gα_s_, β-arrestin2, and downstream ERK signaling, whereas Gα_q/11_ and Gα_12/13_ coupling remain comparatively resistant to pharmacological modulation (Fig. 10G). Setmelanotide provides a particularly clear translational example of this principle: despite being highly effective at driving and, in several mutants, even enhancing canonical Gα_s_ signaling, it cannot universally restore the signaling repertoire lost through disruption of the H8/C-terminal phosphorylation-site region. These findings therefore support a model in which the functional consequence of an MC4R mutation is determined not simply by receptor activity, but by the intersection of three variables receptor sequence, ligand chemotype, and transducer identity. Consequently, therapeutic efficacy cannot be inferred solely from agonist potency or canonical Gα_s_ activation; rather, it depends on whether a ligand can stabilize the receptor conformational states required for the specific signaling pathways that remain structurally accessible. This provides a mechanistic framework for understanding how naturally occurring MC4R variants can generate pathway-selective signaling defects and highlights the potential value of pathway-aware pharmacological profiling in precision MC4R therapeutics.

## 3. Conclusion

The melanocortin-4 receptor (MC4R) is a central regulator of mammalian energy homeostasis (*34*), yet the structural determinants that enable this receptor to engage multiple intracellular signaling pathways have remained incompletely understood. The paradox raised at the outset of this study, how a receptor with such a sparse phosphorylation-site repertoire could achieve the signaling versatility long attributed to an elaborate, multi-site phosphorylation barcode (*3–6*), is directly addressed by our findings. Rather than requiring an extensive phosphorylation code, MC4R achieves broad and pathway-selective signaling control through a small number of strategically positioned residues within helix 8 and the C-terminal tail. In this study, we systematically interrogated these residues and demonstrate that they function not simply as components of a conventional phosphorylation barcode, but as spatially distinct regulators of MC4R signaling architecture. Systematic interrogation of these residues demonstrates that their functional importance extends beyond serving as conventional phosphorylation-dependent regulatory elements; instead, their spatial position of phosphoresidues within the receptor architecture differentially influences receptor activation, transducer engagement, trafficking, and downstream signaling. Notably, helix 8 residue S306, despite having comparatively little prior characterization as a phosphorylation site in the literature, emerged as one of the most consequential residues in the entire panel, producing pronounced effects on receptor activation and transducer engagement. These findings indicate that the functional importance of a regulatory residue is not predicted by the depth of its prior characterization, and that a small, sparse set of regulatory residues, positioned strategically can exert disproportionate control over receptor signaling.

Our results further identify a striking functional distinction between the helix 8 and distal C-terminal regions, both of which have previously been implicated more broadly in MC4R regulation (*16*, *35*). Substitution of helix 8 residues S306 and T312 produced marked alterations in receptor signaling, characterized by enhanced basal Gα_s_-cAMP activity together with impaired β-arrestin2 recruitment and defective downstream signaling and endosomal trafficking. In contrast, mutation of the distal C-terminal residues, particularly S329 and S330, produced a different signaling phenotype, with reduced canonical G-protein signaling and relative preservation of β-arrestin2 recruitment (Fig. 11). These findings indicate that residues positioned within distinct structural regions of MC4R do not contribute equivalently, rather, they differentially regulate the receptor conformations and intracellular interfaces required for pathway-specific signaling (Table. S9). Furthermore, our observation that MC4R preferentially engages β-arrestin2 rather than β-arrestin1 identifies an additional layer of specificity in the receptor’s regulatory and trafficking machinery, consistent with emerging evidence that the two arrestin paralogs adopt distinct, phosphorylation-dependent conformations even when engaging the same receptor (*36*).

**Figure 11:**
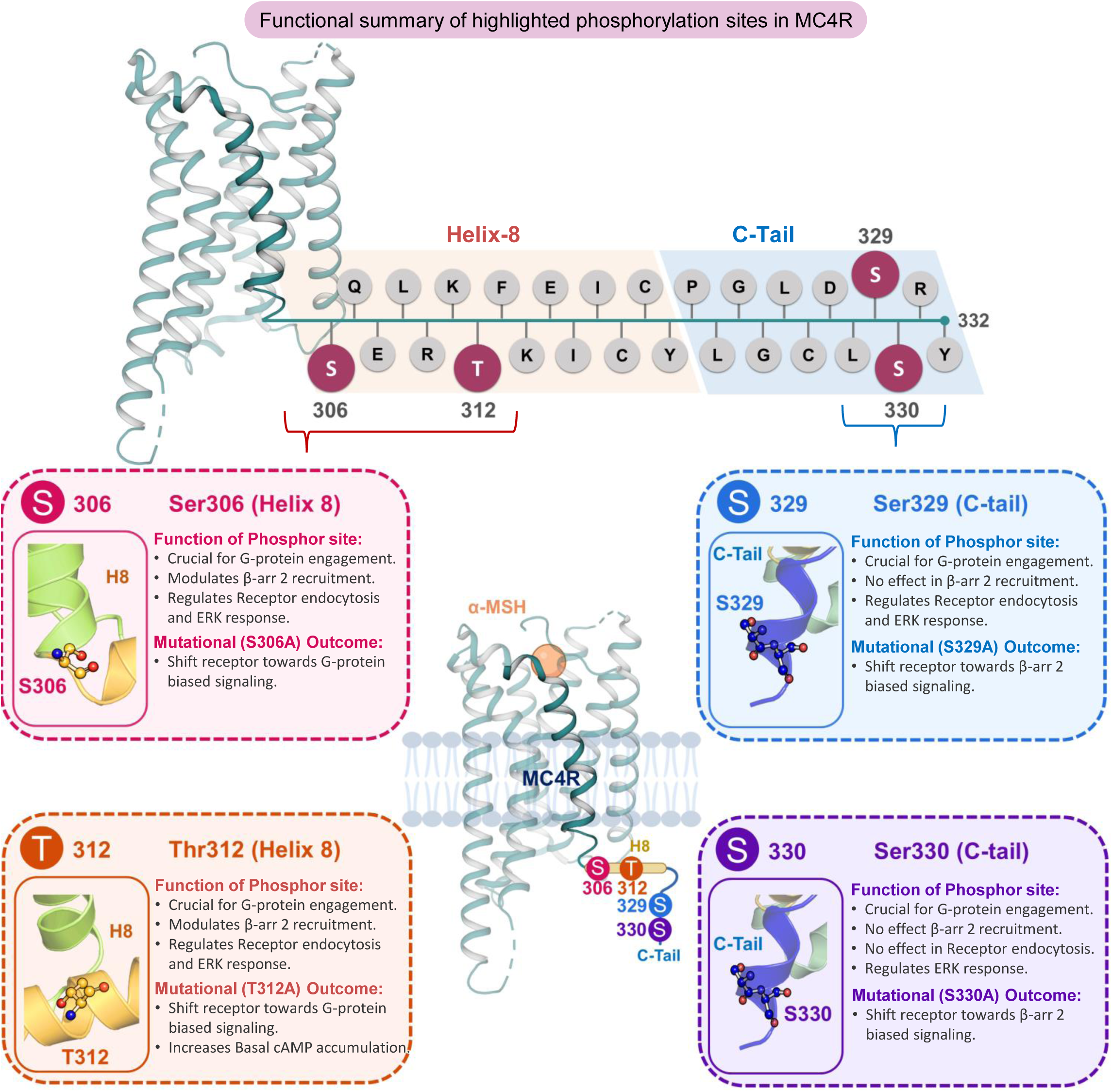
Residue-by-residue functional summary of highlighted phosphorylation sites in MC4R: The upper panel depicts the structural topology and linear amino acid sequence of the proximal Helix 8 and distal C-terminal tail (C-Tail), highlighting the four highly conserved serine and threonine residues investigated in this study. The lower schematic summarizes the distinct functional and pharmacological phenotypes associated with each phosphorylation-deficient alanine mutant upon α-MSH stimulation. Mutations within the proximal Helix 8 domain (S306A and T312A) disrupt β-arrestin 2 recruitment and endosomal trafficking, driving a pronounced G-protein biased signaling profile, with the T312A variant uniquely enhancing basal cAMP accumulation. Conversely, mutations within the distal C-Tail (S329A and S330A) selectively impair G-protein engagement while preserving β-arrestin 2 recruitment, thereby shifting the receptor toward a β-arrestin 2 biased signaling conformation.

Importantly, the consequences of these mutations extended beyond the canonical Gα_s_-β-arrestin axis that has traditionally dominated the MC4R signaling literature (*12*). Across the mutant panel, NFAT-RE-and SRF-RE-dependent responses associated with Gα_q/11_-and Gα_12/13_-mediated signaling, respectively, were markedly attenuated across the receptor variants and remained constrained despite stimulation with chemically diverse agonists. These findings suggest that the helix 8 and C-terminal regions contribute to the structural organization required for productive coupling to multiple transducers, with alternative G-protein pathways displaying substantially greater sensitivity to perturbation than Gα_s_ signaling. Thus, these residues are not simply determinants of phosphorylation-dependent recruitment but form part of a broader structural framework that imposes pathway-specific constraints on MC4R signaling.

A major finding of this study is that these mutation-induced signaling defects are nevertheless pharmacologically plastic. Profiling chemically diverse MC4R agonists, including the superpotent linear and cyclic melanocortin peptides MT-I (*37*) and MT-II (*38*), the clinically studied peptide bremelanotide (*39*), and the structurally distinct small molecule THIQ (*40*); revealed that receptor mutations do not simply eliminate receptor activity; instead, they remodel the relationship between ligand structure, receptor conformation, and transducer engagement. MT-I, MT-II, bremelanotide, and THIQ produced mutation-and pathway-dependent changes in potency and efficacy, and in selected contexts could substantially remodel β-arrestin2 recruitment, Gα_s_ signaling, and downstream ERK responses (Table. S10). The striking reordering of ligand efficacy observed in mutants such as S306A further demonstrates that the functional activity of an agonist is not an invariant property of the ligand but emerges from its interaction with the conformational landscape of the receptor (*41*). Thus, mutation-induced signaling phenotypes can be modulated, and in selected pathways partially bypassed, through ligand-directed pharmacology.

This principle was further highlighted by the clinically approved MC4R agonist setmelanotide (*21*, *22*). Setmelanotide produced a profound enhancement of canonical Gα_s_-cAMP signaling across much of the mutant panel, in some cases producing greater potency and efficacy than MC4R^WT^, consistent with structural evidence that Setmelanotide engages a compact, deep-pocket binding mode that stabilizes a highly active Gα_s_-coupled conformation (*42*), demonstrating that a clinically optimized ligand can exploit residual conformational plasticity within structurally perturbed receptors. However, this pharmacological compensation was not universal. Setmelanotide enhanced potency in β-arrestin2 and ERK signaling without uniformly eliminating variant-specific phenotypes and did not restore the markedly attenuated NFAT-RE-and SRF-RE-dependent responses. Thus, even a highly potent clinical agonist cannot overcome all of the structural constraints imposed by these phosphorylation-site mutations.

Collectively, our findings establish a model in which MC4R signaling is determined by the interplay between receptor sequence, ligand structure, and transducer identity (*41*). MC4R variants therefore should not be viewed simply as receptors that are globally “active” or “inactive.” Rather, individual mutations reshape the conformational ensemble of the receptor, selectively restricting some signaling pathways while preserving or exposing others, and different agonists can differentially exploit this altered signaling landscape. Together with the phenotype of S306, these results directly answer the question posed at the outset of this study: MC4R does not require an extensive, multi-site phosphorylation barcode to achieve broad signaling versatility, since a sparse set of phosphorylation-site residues, including previously uncharacterized ones, is sufficient to encode pathway-selective control. This provides a mechanistic framework for understanding how naturally occurring MC4R variants can produce pathway-selective functional phenotypes and why their pharmacological behavior cannot be predicted solely from canonical Gα_s_ activity. More broadly, these findings highlight the value of pathway-aware pharmacological profiling and ligand-directed receptor modulation for understanding, and potentially therapeutically exploiting, signaling-deficient MC4R variants associated with metabolic disease.

## 4. Material and Methods

### Reagents and Plasmids

The wild-type human MC4R (hMC4R), hMC4R-smBiT, LgBiT-β-arrestin 1 and 2, and LgBiT-FYVE constructs, all inserted into the pcDNA3.1(+) mammalian expression vector, were synthesized and procured from GenScript and Twist Bioscience. Site-directed mutagenesis to introduce specific alanine substitutions at the phosphorylation sites within helix 8 and the C-terminal region of the hMC4R construct was performed in-house utilizing the Q5® Site-Directed Mutagenesis Kit (New England Biolabs, cat. no. E0554S). The corresponding phosphorylation site mutations within the hMC4R-smBiT constructs were custom-synthesized by Twist Bioscience.

### Site-Directed Mutagenesis of MC4R Constructs

To systematically interrogate the structural determinants located within the proximal helix 8 and distal c-terminal tail of the human MC4R, we have targeted highly conserved serine (S) and threonine (T) phosphorylation sites for alanine (A) substitution. Custom oligonucleotide primers encoding the specific nucleotide mismatches were designed and commercially synthesized by Sigma-Aldrich; while mutagenesis was executed utilizing the Q5^®^ Site-Directed Mutagenesis Kit (New England Biolabs, cat. no. E0554S) in strict accordance with the manufacturer’s recommended protocols for exponential amplification, kinase/ligase/DpnI (KLD) treatment, and subsequent transformation. This approach successfully yielded four single-point mutant expression constructs MC4R^S306A^, MC4R^T312A^, MC4R^S329A^, and MC4R^S330A^ as well as one double-mutant construct, MC4R^S329A:S330A^. Prior to their application in cellular and functional assays, the successful integration of all targeted mutations, alongside the absence of off-target sequence alterations within the receptor reading frame, was rigorously validated for all generated plasmids via Sanger sequencing performed by Macrogen, South Korea and IISER, Bhopal.

### Cell Culture

Human Embryonic Kidney (HEK) 293T cells were obtained from the National Centre for Cell Science (NCCS, Pune, India). Cells were maintained in Dulbecco’s Modified Eagle Medium (DMEM; Gibco, cat. no. 12800017), supplemented with 10% (v/v) heat-inactivated fetal bovine serum (FBS; Gibco, cat. no. 10270106) and 1% Penicillin-Streptomycin (10,000 U/mL; Gibco, cat. no. 15140122). Cultures were maintained in 100 mm sterile petri dishes (Abdos) in a humidified incubator at 37°C with 5% CO_2_. Cells were routinely passaged upon reaching 70–80% confluency using 0.25% Trypsin-EDTA (Gibco, cat. no. 25200072) for cellular detachment.

### Reporter Systems

For functional signaling assays, the pGloSensor™-22F cAMP plasmid (cat. no. E2301) was utilized for real-time cyclic AMP monitoring. Downstream transcriptional activity was evaluated using specific luciferase reporter plasmids: pGL4.30[luc2P/NFAT-RE/Hygro] (cat. no. E8481) for Gα_q/11_ signaling, pGL4.34[luc2P/SRF-RE/Hygro] (cat. no. E1350) for Gα_12/13_ signaling, and pGL4.33[luc2P/SRE/Hygro] (cat. no. E1340) for MAPK/ERK signaling. All reporter plasmids were purchased from Promega.

### Pharmacological Agents

The endogenous neuropeptide α-MSH (cat. no. M4135) was obtained from Sigma-Aldrich. The synthetic pharmacological panel, which included Melanotan I (MT-I; MedChemExpress, cat. no. HYN2466), Melanotan II (MT-II; MedChemExpress, cat. no. HYP0267), Bremelanotide (MedChemExpress, cat. no. SML2756), the small-molecule agonist THIQ (Sigma-Aldrich, cat. no. SML2449), and the FDA-approved therapeutic Setmelanotide (MedChemExpress, cat. no. HY-19870), were all purchased.

All other laboratory chemicals, buffers, and reagents were of standard analytical or molecular biology grade and were purchased from Sigma-Aldrich; Gibco or standard commercial suppliers unless otherwise stated.

### Quantification of Receptor Surface Expression via Whole-Cell ELISA

To evaluate and normalize the cell-surface expression of MC4R^WT^ and its corresponding phosphorylation site variants MC4R^S306A^, MC4R^T312A^, MC4R^S329A^, MC4R^S330A^, and MC4R^S329A:S330A^; we performed a colorimetric whole-cell enzyme-linked immunosorbent assay (ELISA) targeting the N-terminal FLAG epitope of the receptor constructs (*43*). HEK293T cells were transiently transfected with the respective receptor plasmids utilizing Polyethylenimine (PEI; 1 mg/mL) at a DNA-to-PEI ratio of 1:3. Post-transfection, the cells were seeded at a density of 0.2 million cells/well into 24-well plates pre-coated with 0.01% poly-D-lysine (Sigma-Aldrich, cat. no. P0899). After a 24-h incubation, the plates were transferred to ice, and the cells were fixed using 4% (w/v) paraformaldehyde (pH 6.9) for 20 min. Following fixation, the cells were washed three times with 1× Tris-buffered saline (TBS; 150 mM NaCl, 50 mM Tris-HCl, pH 7.4). To prevent non-specific antibody binding, cells were then incubated in a blocking buffer consisting of 1% (w/v) bovine serum albumin (BSA) (SRL, cat. no. 8517) in 1× TBS for 90 min. Cells were subsequently incubated for 90 min with a horseradish peroxidase (HRP)-conjugated anti-FLAG M2 antibody (1:5000 dilution; Sigma-Aldrich, cat. no. A8592) prepared in 1% BSA. Unbound antibodies were removed through three consecutive washes with 0.5% BSA in TBS. For colorimetric signal development, 200 ul of tetramethylbenzidine (TMB) ELISA substrate (Thermo Fisher Scientific, cat. no. 34028) was added to each well. Upon the visual appearance of a blue reaction product, the enzymatic reaction was terminated by transferring 100 uL of the developed solution into a fresh 96-well plate pre-loaded with 100 uL of 1 M H_2_SO_4_. The absorbance of the final solution was immediately recorded at 450 nm using a multimode microplate reader (BMG Labtech). To accurately normalize the surface expression signals against variations in cell seeding density across different wells, we performed a Janus Green cellular staining protocol. The remaining TMB solution in the 24-well plate was aspirated, and the cells were washed once with 1× TBS prior to incubation with 0.2% (w/v) Janus Green stain (Sigma-Aldrich, cat. no. 201677) for 20 min. Following four extensive washes with distilled water to remove excess dye, the incorporated stain was eluted by adding 800 uL of 0.8 N HCl per well. A 200 uL aliquot of this eluate was transferred to a 96-well plate, and the absorbance was measured at 595 nm. The final, normalized surface expression for each receptor construct was calculated as the ratio of the antibody-specific signal (*OD_450_*) to the Janus Green cell density signal (*OD_595_*).

### cAMP Accumulation via GloSensor Assay

Ligand-induced Gα_s_ activation was quantified by monitoring real-time intracellular cyclic AMP (cAMP) accumulation using the live-cell GloSensor assay, adapting previously described methodologies (*44*). Briefly, HEK293T cells were transiently co-transfected with 4 ug of the respective FLAG-tagged receptor construct and 3 ug of the pGloSensor™-22F cAMP plasmid. At 14-16 h post-transfection, cells were harvested and resuspended in assay buffer (1× HBSS, pH 7.4, supplemented with 20 mM HEPES) containing 1 mg/mL D-luciferin (GoldBio, cat. no. LUCNA-1G). Cells were seeded into 96-well white microplates (SPL Life Sciences, cat. no. 30169) at a density of 1.25 x 10^5^ cells/100uL per well. The plates were incubated at 37 °C for 90 min to allow substrate equilibration, followed by an additional 30 min incubation at room temperature. For receptor stimulation, serial dilutions of the pharmacological agents (ranging from 1 pM to 1 uM) were prepared and added to the respective wells. Luminescence was immediately recorded using a multimode microplate reader (BMG Labtech). Data were normalized by setting the maximal agonist-induced response to 100%, and concentration-response curves were plotted and analyzed using GraphPad Prism 10.

### NanoBiT-Based β-Arrestin Recruitment Assay

Agonist-induced β-arrestin recruitment was evaluated utilizing the NanoLuc Binary Technology (NanoBiT) complementation assay, following established protocols (*43*). HEK293T cells were transiently co-transfected with 3.5 ug of either MC4R^WT^-smBiT or the specified MC4R^Mutant^-smBiT constructs (bearing an N-terminal FLAG-tag and a C-terminal smBiT tag), alongside 3.5 ug of either LgBiT-βarr1 or LgBiT-βarr2 (harboring an N-terminal LgBiT tag). At 14–16 h post-transfection, cells were washed once with 1× PBS, detached via trypsinization, and seeded into 96-well white microplates at a density of 1 x 10^5^ cells/well. The cells were suspended in assay buffer containing 1× HBSS, 5 mM HEPES (pH 7.4), 0.01% (w/v) BSA, and 10 uM coelenterazine native (GoldBio, cat. no. CZ05). Plates were incubated at 37 °C for 90 min, followed by 30 min at room temperature. Following ligand stimulation, bioluminescence was recorded using a multimode plate reader. The maximal agonist concentration response was normalized to 100%, and analysis was performed via GraphPad Prism 10.

### NanoBiT-Based Receptor Endocytosis Assay

To dynamically monitor the agonist-induced internalization of MC4R constructs into early endosomes, we employed a FYVE-domain-based NanoBiT trafficking assay (*43*). HEK293T cells were co-transfected with 2 ug of the respective smBiT-tagged receptor construct (MC4R^WT^-smBiT or MC4R^Mutant^-smBiT) and 5 ug of the early endosome marker LgBiT-FYVE. At 14–16 h post-transfection, cells were washed, trypsinized, and seeded into 96-well microplates (1 x 10^5^ cells/well) in the same coelenterazine-supplemented assay buffer described for the β-arrestin recruitment assay. After the standard equilibration periods (90 min at 37 °C, 30 min at room temperature), baseline luminescence was established prior to ligand addition. Real-time endosomal accumulation was recorded continuously on a multimode plate reader. Data normalization was done with respect to time as per the cycle and graphical visualization at the end was conducted using GraphPad Prism 10.

### Immunoblotting for ERK1/2 Phosphorylation

To evaluate the rapid, G-protein-dependent phase of ERK1/2 activation across the receptor variants, agonist-induced ERK1/2 phosphorylation was quantified via Western blotting (*2*). HEK293T cells at 60– 70% confluency were transfected with 4 µg of the MC4R(WT) construct or 5 µg of the respective mutant constructs (MC4R^S306A^, MC4R^T312A^, MC4R^S329A^, MC4R^S330A^, or MC4R^S329A:S330A^). This differential plasmid loading was utilized to ensure comparable baseline surface expression across all variants. At 24 h post-transfection, cells were harvested and seeded into 6-well tissue culture plates at a density of 1 × 10^6^ cells per well. The following day, the growth media was removed, and the cells were subjected to serum starvation in serum-free DMEM for 12 h to minimize basal kinase activity. Cells were then subsequently stimulated with 0.1 µM and 1 µM of the endogenous agonist α-MSH for exactly 5 min. Immediately following stimulation, the media was rapidly aspirated, and cellular signaling was quenched by lysing the cells directly in 100 µL of 2x SDS protein loading buffer. Cellular lysates were denatured by heating at 100°C for 15 min and clarified by centrifugation at 14,800 rpm for 15–20 mins. Then an equal volume (15 µL) of the clarified lysates were loaded per well, resolved via 12% SDS-polyacrylamide gel electrophoresis (SDS-PAGE), and transferred to a PVDF membrane. To detect active kinase levels, the membranes were immunoblotted with a primary anti-phospho-ERK1/2 antibody (1:5000 dilution; Cell Signaling Technology, cat. no. 9101). To normalize the phosphorylation signal against the overall target protein content, the blots were subsequently evaluated using an anti-total-ERK1/2 primary antibody (1:5000 dilution; Cell Signaling Technology, cat. no. 9102). Because the primary antibodies were unconjugated, a horseradish peroxidase (HRP)-conjugated anti-rabbit IgG (1:5000 dilution; Sigma-Aldrich, cat. no. A9169) was utilized as the secondary detection antibody. The chemiluminescent signals were captured using a ChemiDoc Imaging System (Bio-Rad), and precise densitometric quantification of the protein bands was performed utilizing Image Lab 6.1 software (Bio-Rad).

### ERK Signaling via SRE Reporter Assay

Terminal MAPK/ERK signaling was assessed utilizing a Serum Response Element (SRE) luciferase reporter assay, according to previously outlined methods (*45*). HEK293T cells were transiently co-transfected with a mixture of 3.5 ug of the relevant FLAG-tagged receptor construct and 3.5 ug of the pGL4.33[luc2P/SRE/Hygro] reporter plasmid. At 14–16 h post-transfection, cells were harvested and seeded into 96-well plates (1 x 10^5^ cells/well) in complete growth media. After an 8 h attachment period, the complete media was carefully aspirated, and the cells were subjected to serum starvation overnight at 37°C in serum-free DMEM to minimize basal kinase activity. The following morning, the cells were stimulated with designated concentrations of pharmacological agents for 6 h. Post-stimulation, the ligand-containing media was aspirated, and 100 uL of detection buffer (1× HBSS, 20 mM HEPES, pH 7.4, supplemented with 0.5 mg/mL D-luciferin) was added per well. Luminescence was quantified immediately. The data were normalized against the maximal agonist-induced response and analyzed using GraphPad Prism 10.

### Evaluation of Gα_q/11_ and Gα_12/13_ Signaling

Receptor-mediated coupling to the Gα_q/11_ and Gα_12/13_ pathways was evaluated using Nuclear Factor of Activated T-cells (NFAT) and Serum Response Factor-Response Element (SRF-RE) luciferase reporter assays, respectively (*45*). In parallel experimental setups, HEK293T cells were co-transfected with 3.5 ug of the FLAG-tagged MC4R constructs and 3.5 ug of either the pGL4.30[luc2P/NFAT-RE/Hygro] plasmid (for activation of Gα_q/11_ detection) or the pGL4.34[luc2P/SRF-RE/Hygro] plasmid (for activation of Gα_12/13_ detection). At 14–16 h post-transfection, cells were seeded into 96-well plates (1 x 10^5^ cells/well) in complete media. Consistent with the SRE assay protocol, the cells were allowed to adhere for 8 h before the media was replaced with serum-free DMEM for overnight starvation at 37 °C. The cells were subsequently stimulated with the respective ligands for 6 h. The media was then replaced with 100 uL of D-luciferin-supplemented assay buffer, and the resultant luminescence was measured immediately. All data were normalized to the maximal agonist-induced signal (100%) and analyzed using GraphPad Prism 10.

### Quantifying bias factor

Two important pharmacological parameters for bias calculation obtained from concentration-response curves are E_max_ and EC_50_. By measuring cellular responses with serially diluted ligands and fitting the concentration response to a sigmoidal curve, E_max_ and EC_50_ are calculated. Emax represents the maximum response subtracted from the minimal response. EC_50_ indicates the concentration of an agonist that gives half of the E_max_ response. Whilst agonists can be classified as full or partial agonists based on their efficacy, the classification is assay system-dependent since the degree of signal amplification varies depending on the assay system. (*46*)

Amongst various ways to evaluate bias, the simplest and most practical way is to calculate a ratio of the E_max_ and EC_50_ value of each ligand. Another commonly used method is the Black-Leff operational model, which evaluates bias by calculating ΔΔLog (τ/KA) (*47–50*). This model was built to formulate pharmacological agonism, taking into account the formation process of the agonist receptor-signal transducer complex (*49*). In this model, τ /K_A_, referred to as the transduction coefficient, is composed of the transducer ratio (τ), which is dictated by the receptor density and coupling efficiency, and the agonist receptor dissociation constant (K_A_). Further details of this approach are described elsewhere (*47–50*). The relative activity of the test ligand to the reference ligand must be calculated for each assay. The relative activity to the reference ligand is calculated by the following equation:

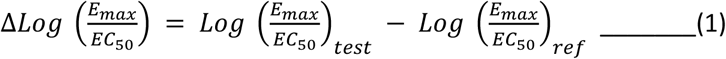

Equation (1) is equivalent to the logarithmic value of the intrinsic relative activity (RAi), which is used to compare the activities of ligands in the same assay (*51*).

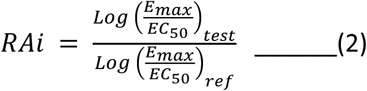

These evaluations encompass both the efficacy and potency of the ligand. To assess bias activity, the Δ Log (Emax/EC50) value of one pathway is subtracted from that of another pathway, yielding ΔΔ Log (Emax/EC50) value, as shown in the following equation.

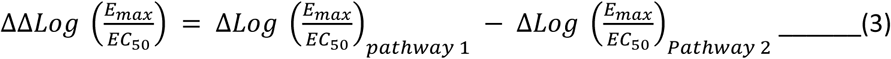

Bias factor, which indicates the fold change in biased signaling activity between two pathways, can be obtained as the inverse logarithm of ΔΔ Log (Emax/EC50).

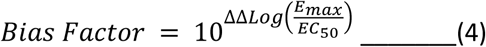

Bias factor or ΔΔ Log (Emax/EC50) value was then visualized in XY scatter plot using GraphPad Prism 10 software.

## Author contributions

M.P.B. performed almost all the experiments with some assistance from other authors. S.A. assisted with the β-arrestin recruitment assays. M.A. contributed to data analysis and figure preparation. N.S. assisted with selected experiments and experimental procedures. M.B. and P.K. conceptualized and designed the study, supervised the research, and contributed to manuscript preparation, editing, and revision. All authors reviewed and approved the final manuscript.

## Acknowledgement

M.B.’s lab is supported by the Early Career Fellowship from the India Alliance Wellcome Trust (IA/E/20/1/505691), Institute Seed Grant-IIT Jammu (SGT-100081), SERB-SRG Grant (SRG/2023/000424) and ICMR Grant (IIRP-2023-2121). P.K.’s lab is supported by IISER Bhopal Start-up Grant (IISERB/R&D/2024-25/66) and ANRF (ANRF/ECRG/2024/000398/LS).

## Data, code, and materials availability

All data needed to evaluate the conclusions in the paper are present in the paper or the Supplementary Materials. All plasmids and materials generated in this study will be distributed upon reasonable request and completion of a materials transfer agreement with IIT Jammu.

## SUPPLEMENTARY FIGURES

**Figure S1:**
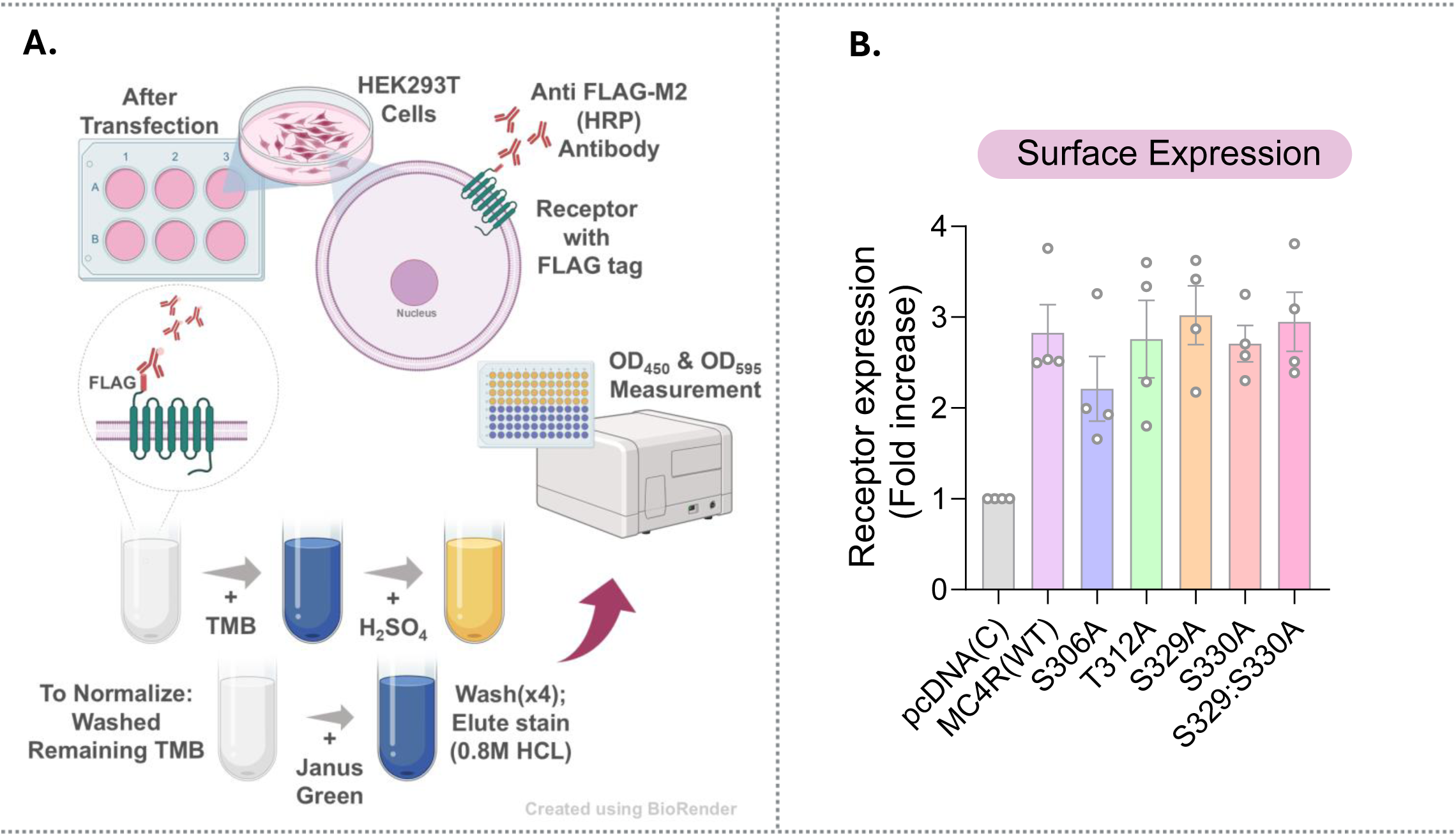
Surface Expression of MC4R Phosphorylation Site Mutants: **A.** Schematic representation of colorimetric whole-cell enzyme-linked immunosorbent assay (ELISA) targeting the N-terminal FLAG epitope of the receptor constructs. **B.** Surface expression in bar graph, n=4, was analyzed using one-way ANOVA with Dunnett test where no significant differences seen in expression; error bar indicates ±SEM.

**Figure S2:**
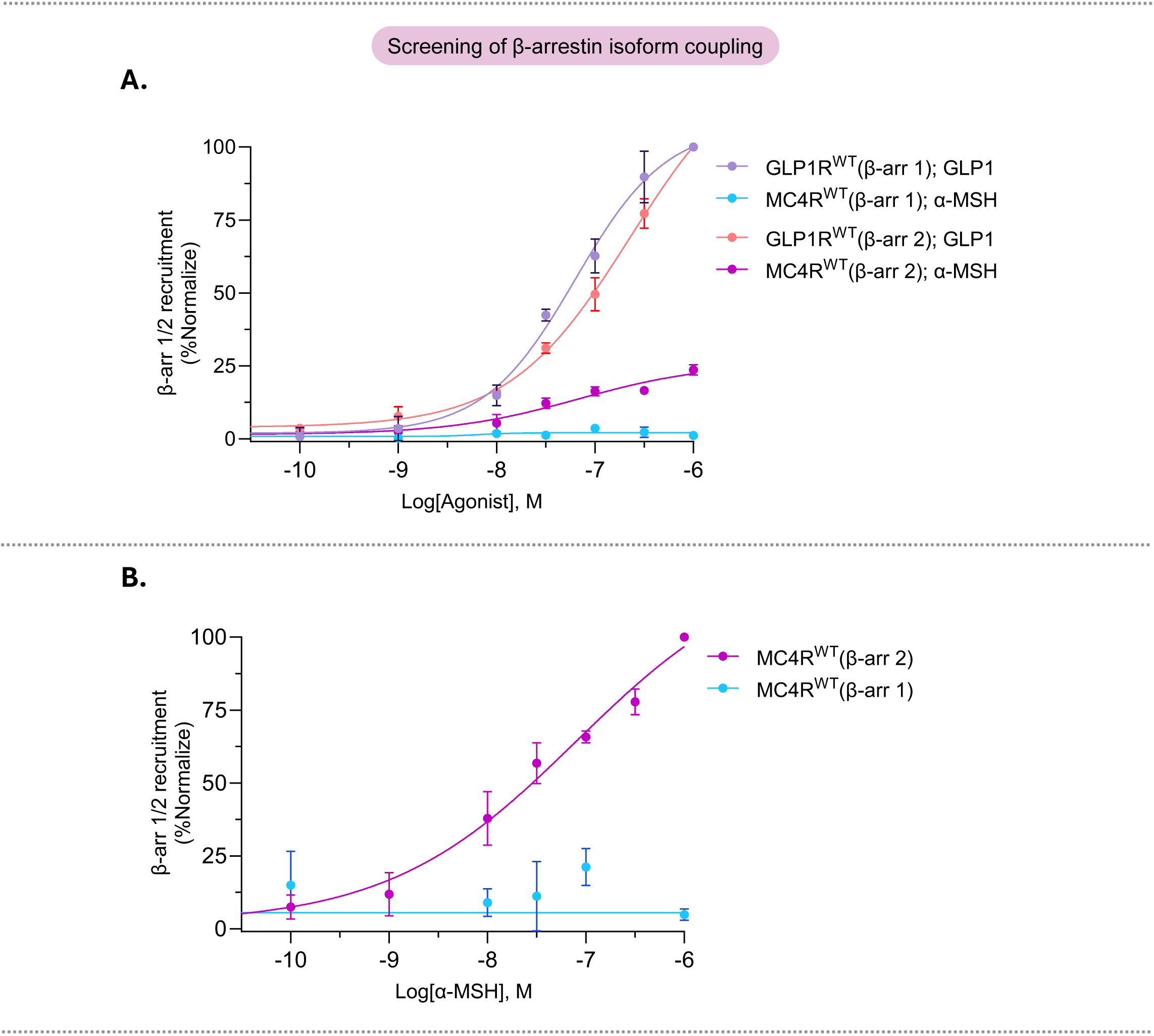
Screening of β-arrestin 1 & 2 recruitment. **A.** Using NanoBiT based assay, MC4R^WT^ along with GLP1R^WT^ (as positive control) were both tested for β-arrestin 1/2 recruitment along with their respective endogenous ligand (α-MSH and GLP1); n=3; **B.** MC4R^WT^ compared for β-arrestin isoform coupling (n=3); % normalized of luminescence data in HEK293T cells; error bar indicates ±SEM.

**Figure S3:**
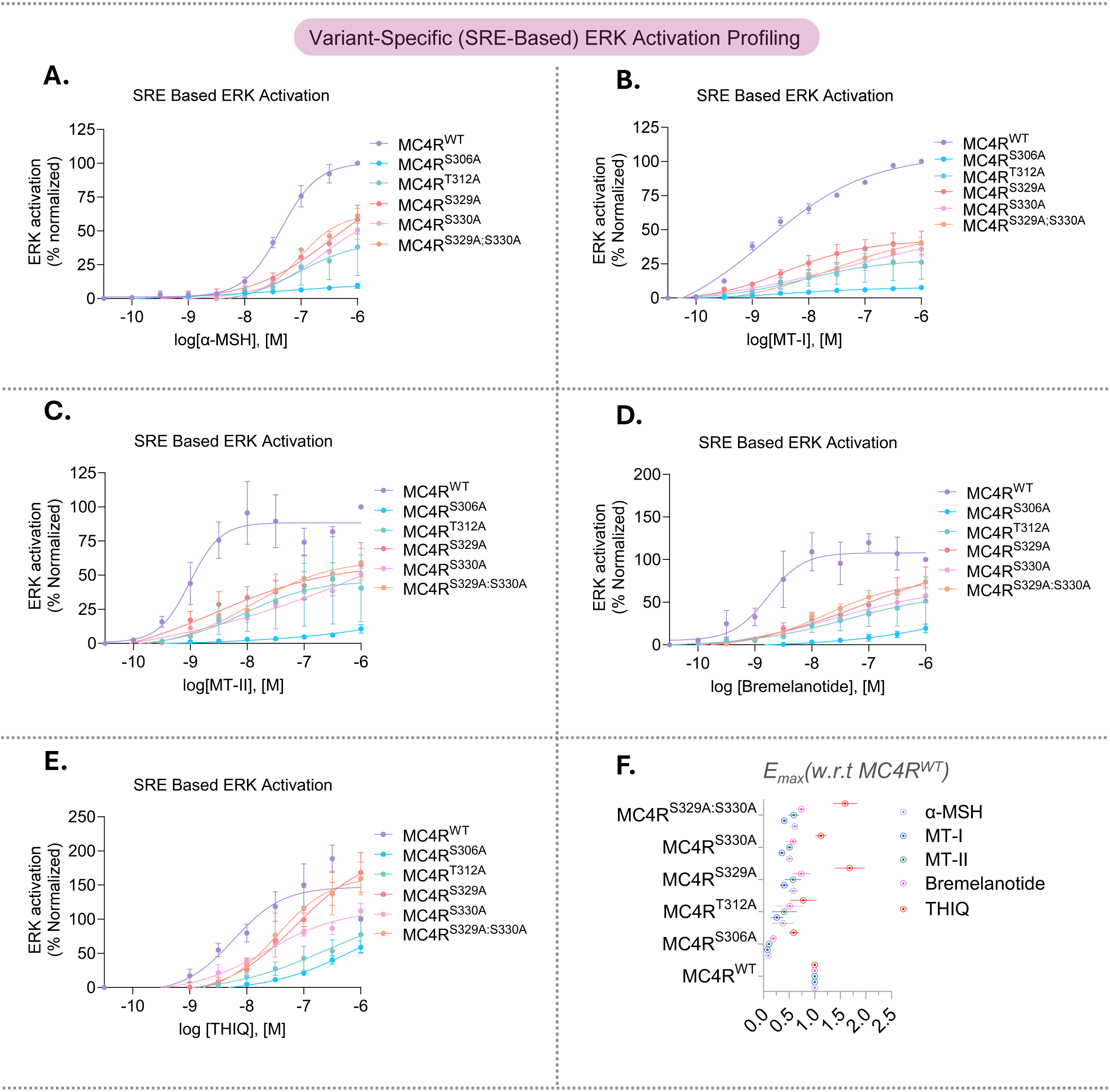
Variant specific SRE based ERK activation Profiling with its Pharmacological agents: Comparison of MC4R variants, with a panel of agonist **A.** α-MSH**; B.** MT-I**; C.** MT-II**; D.** BREMELANOTIDE**; E.** THIQ; to check ERK activation via luminescence-based SRE reporter assay; n=4; % normalized of luminescence data in HEK293T cells**; F.** Shows the Emax variability while comparing with the variant against the set of agonist; error bar indicates ±SEM.

**Figure S4:**
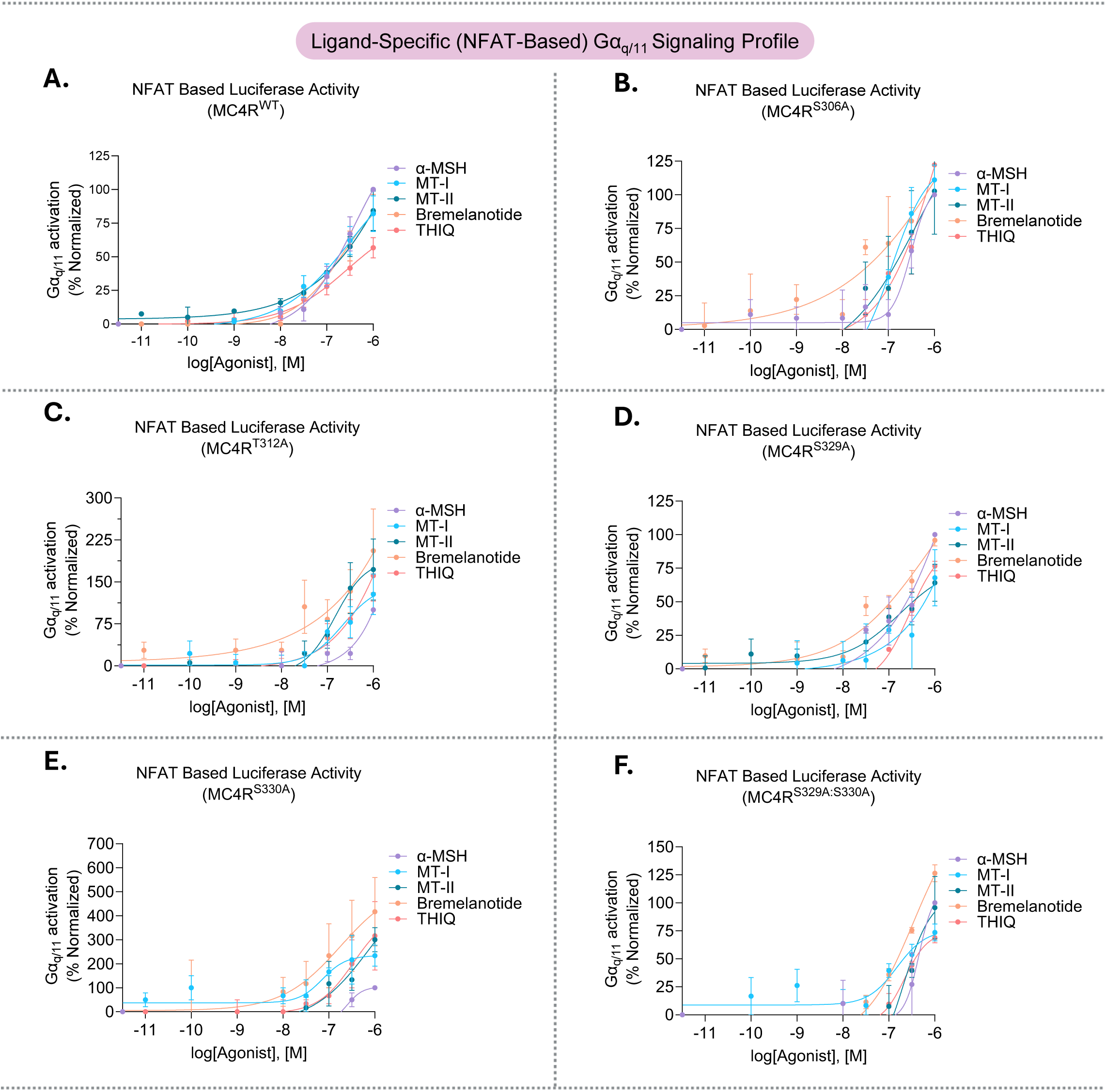
Ligand-Specific Gαq/11 Profiling with MC4R Variants: Comparison with MC4R variants, **A.** MC4R^WT^**; B.** MC4R^S306A^**; C.** MC4R^T312A^**; D.** MC4R^S329A^**; E.** MC4R^S330A^**; F.** MC4R^S329A:S330A^; and set of pharmacological agents to check Gαq/11 activation via luminescence-based NFAT-RE reporter assay; n=3; % normalized of luminescence data in HEK293T cells; error bar indicates ±SEM.

**Figure S5:**
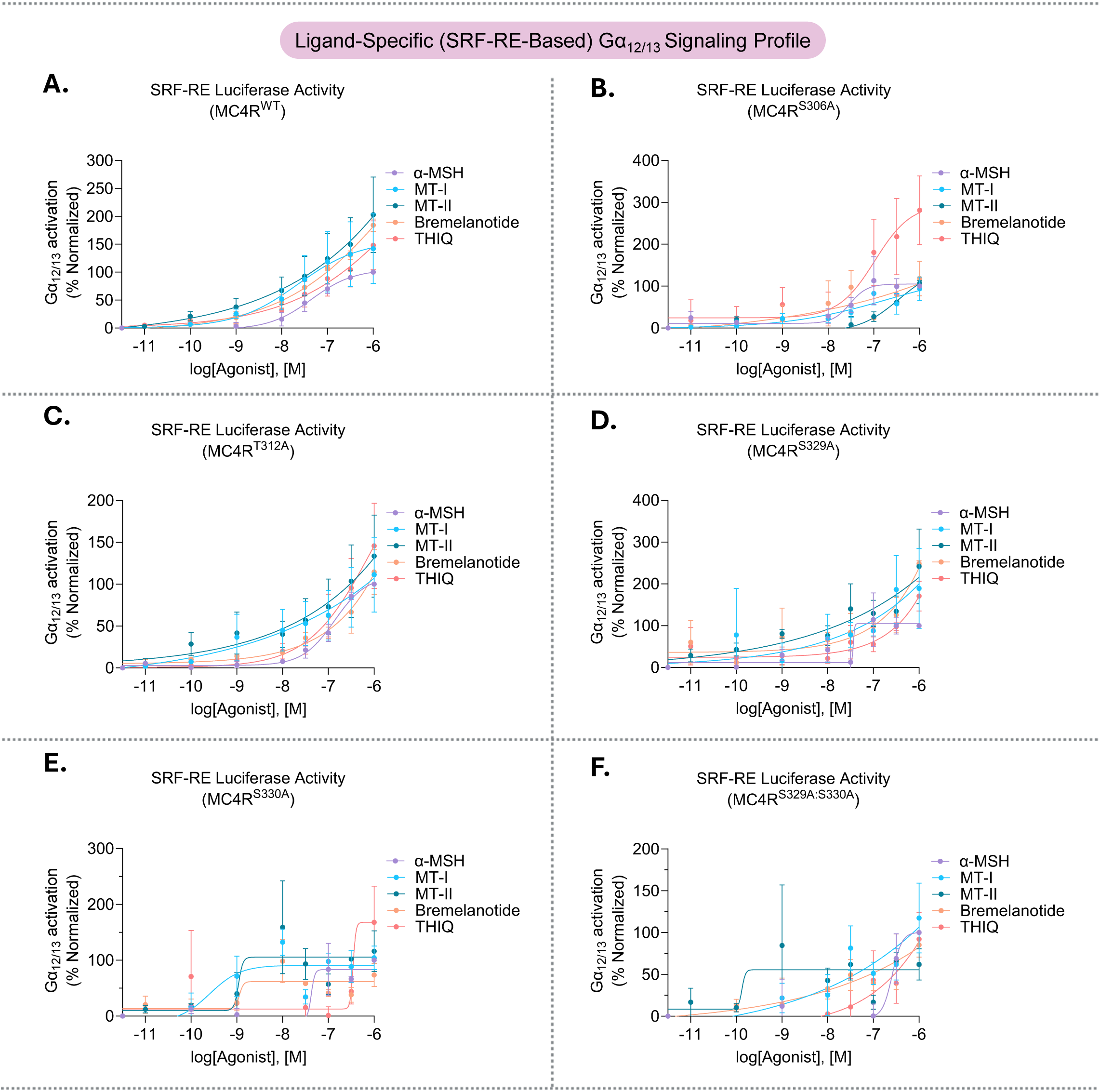
Ligand-Specific Gα12/13 Profiling with MC4R Variants: Comparison with MC4R variants, **A.** MC4R^WT^**; B.** MC4R^S306A^**; C.** MC4R^T312A^**; D.** MC4R^S329A^**; E.** MC4R^S330A^**; F.** MC4R^S329A:S330A^; and set of pharmacological agents to check Gα12/13 activation via luminescence-based SRF-RE reporter assay; n=3; % normalized of luminescence data in HEK293T cells; error bar indicates ±SEM.

**Figure S6:**
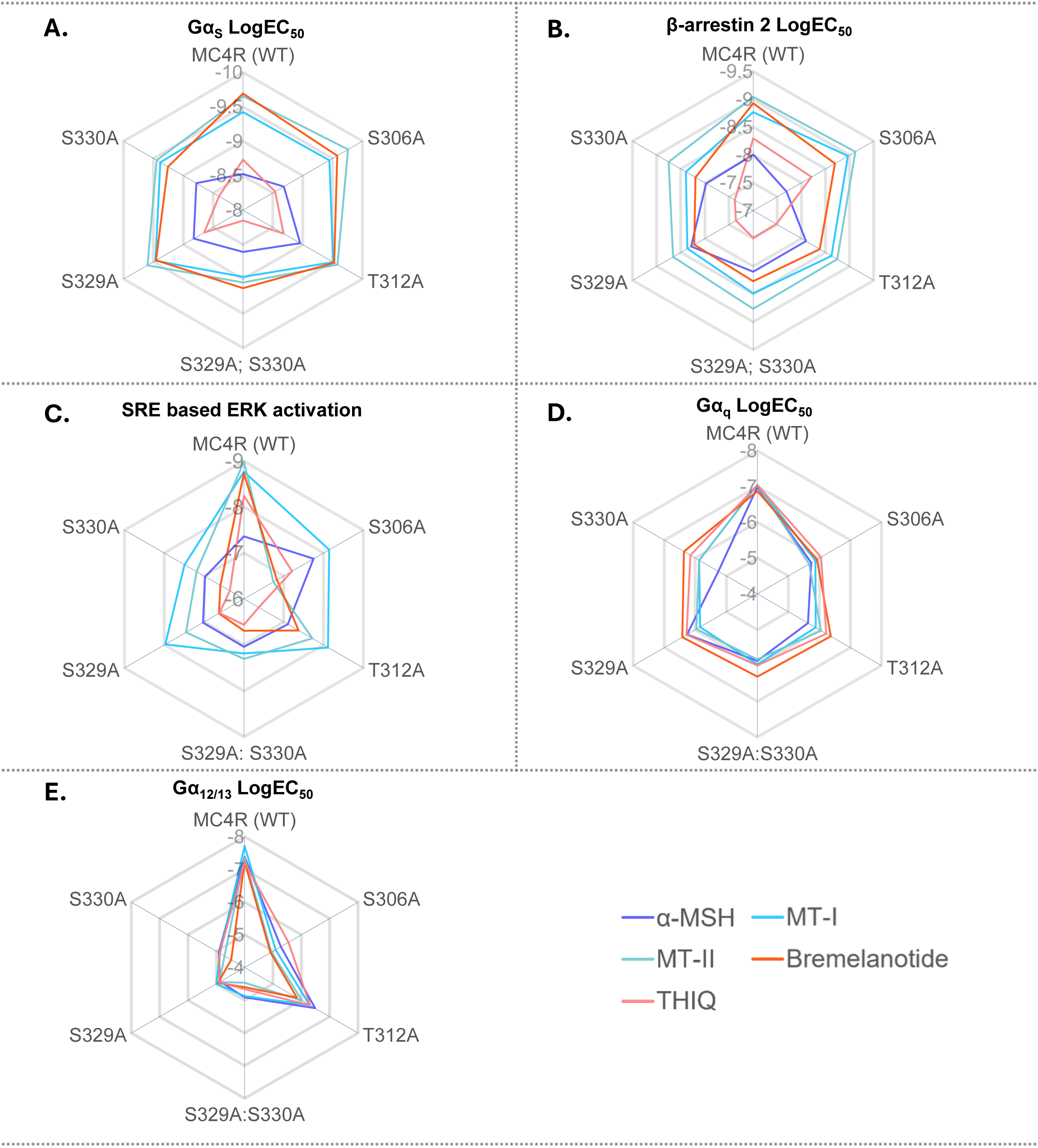
Radar Plot: Depicts the LogEC_50_ values of **A.** Gα_s_ activation profiling via luminescence-based cAMP accumulation assay**; B.** β-arrestin 2 recruitment via NanoBiT based assay**; C.** ERK activation via SRE Reporter luminescence-based assay**; D.** Gα_q_ activation via NFAT-RE Reporter luminescence-based assay**; E.** Gα_12/13_ activation via SRF-RE Reporter luminescence-based assay; in HEK293T cells; simultaneously for each pharmacological agent profile with each of the variants.

---

**Table S1.** Calculation of receptor mediated bias factor with its endogenous ligand α-MSH.

| Receptor | G protein (Gα<sub>s</sub>) |  |  |  | β-arrestin 2 |  |  |  | Receptor bias |  |
| --- | --- | --- | --- | --- | --- | --- | --- | --- | --- | --- |
|  | Emax (%)<sup>*</sup> | EC<sub>50</sub> (nM)<sup>#</sup> | Log (Emax/EC<sub>50</sub>) | ΔLog (Emax/EC<sub>50</sub>) | Emax (%)<sup>*</sup> | EC<sub>50</sub> (nM)<sup>#</sup> | Log (Emax/EC<sub>50</sub>) | ΔLog (Emax/EC<sub>50</sub>) | ΔΔLog (Emax/EC<sub>50</sub>) | Biased |
| MC4R (WT) | 100 | 1.94536 | 8.7 | 0.0 | 100 | 11.3762729 | 7.9 | 0.0 | 0.0 | Balanced |
| S306A | 58.212148 | 1.51008 | 8.6 | -0.1 | 26.58701 | 136144.468 | 3.3 | -4.7 | -4.5 | G protein |
| T312A | 62.806326 | 0.69343 | 9.0 | 0.2 | 48.707147 | 814.704284 | 5.8 | -2.2 | -2.4 | G protein |
| S329A | 53.70534 | 2.55859 | 8.3 | -0.4 | 111.26003 | 8.91250938 | 8.1 | 0.2 | 0.5 | Arrestin |
| S330A | 62.446928 | 2.37684 | 8.4 | -0.3 | 118.7396 | 6.30957344 | 8.3 | 0.3 | 0.6 | Arrestin |
| S329A:S330A | 55.321652 | 2.20293 | 8.4 | -0.3 | 109.1218 | 8.79022517 | 8.1 | 0.1 | 0.5 | Arrestin |

**Table S2.** Calculation of receptor mediated bias factor with its FDA approved ligand Setmelanotide.

| Receptor | G protein (Gα<sub>s</sub>) |  |  |  | β-arrestin 2 |  |  |  | Receptor bias |  |
| --- | --- | --- | --- | --- | --- | --- | --- | --- | --- | --- |
|  | Emax (%)<sup>*</sup> | EC<sub>50</sub> (nM)<sup>#</sup> | Log (Emax/EC<sub>50</sub>) | ΔLog (Emax/EC<sub>50</sub>) | Emax (%)<sup>*</sup> | EC<sub>50</sub> (nM)<sup>#</sup> | Log (Emax/EC<sub>50</sub>) | ΔLog (Emax/EC<sub>50</sub>) | ΔΔLog (Emax/EC<sub>50</sub>) | Biased |
| MC4R (WT) | 100 | 0.069183097 | 10.2 | 0.0 | 100 | 0.603948629 | 9.2 | 0.0 | 0.0 | Balanced |
| S306A | 99.789 | 0.036307805 | 10.4 | 0.3 | 11.800917 | 5188.000389 | 4.4 | -4.9 | -5.1 | G protein |
| T312A | 78.07163333 | 0.030902954 | 10.4 | 0.2 | 48.81958 | 192.3091729 | 6.4 | -2.8 | -3.1 | G protein |
| S329A | 136.7967 | 0.022908677 | 10.8 | 0.6 | 86.15555667 | 0.895364766 | 9.0 | -0.2 | -0.9 | Arrestin |
| S330A | 124.4551 | 70.9577768 | 7.2 | -2.9 | 97.76546667 | 0.393550075 | 9.4 | 0.2 | 3.1 | Arrestin |
| S329A:S330A | 36.48813333 | 0.016982437 | 10.3 | 0.2 | 112.3543667 | 1.721868575 | 8.8 | -0.4 | -0.6 | Arrestin |

**Table S3.**
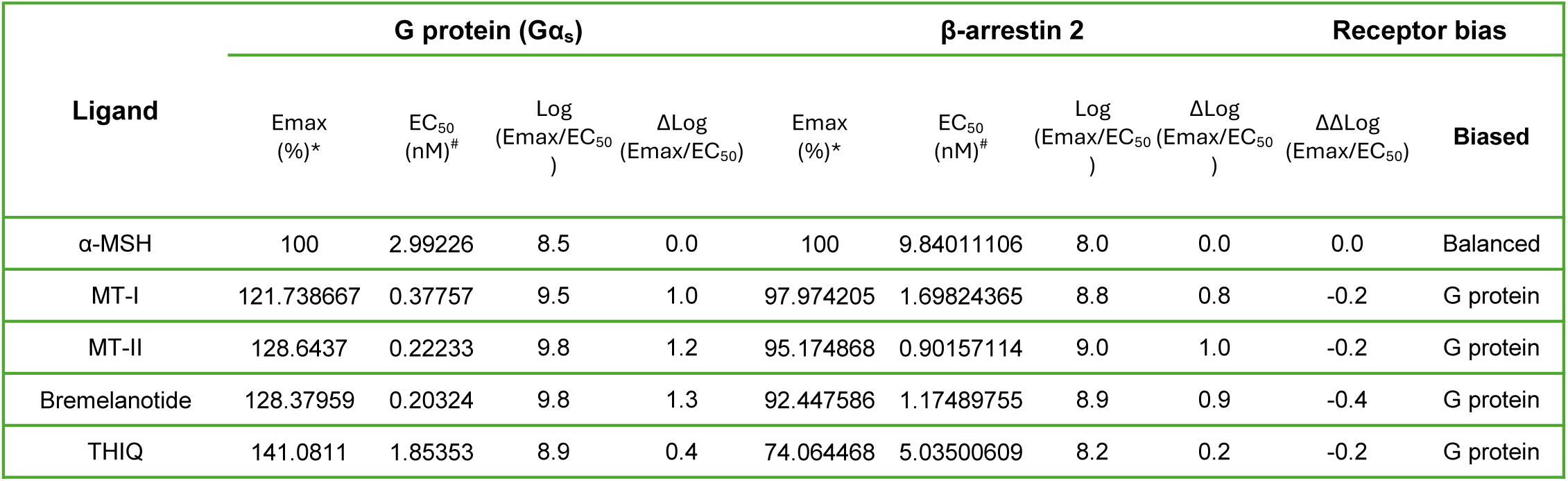
Calculation of ligand mediated bias factor with MC4R^WT^ receptor.

**Table S4.** Calculation of ligand mediated bias factor with MC4R^S306A^ variant.

| Ligand | G protein (Gα<sub>s</sub>) |  |  |  | β-arrestin 2 |  |  |  | Receptor bias |  |
| --- | --- | --- | --- | --- | --- | --- | --- | --- | --- | --- |
|  | Emax (%)<sup>*</sup> | EC<sub>50</sub> (nM)<sup>#</sup> | Log (Emax/EC<sub>50</sub>) | ΔLog (Emax/EC<sub>50</sub>) | Emax (%)<sup>*</sup> | EC<sub>50</sub> (nM)<sup>#</sup> | Log (Emax/EC<sub>50</sub>) | ΔLog (Emax/EC<sub>50</sub>) | ΔΔLog (Emax/EC<sub>50</sub>) | Biased |
| α-MSH | 100 | 2.07491 | 8.7 | 0.0 | 100 | 20.4173794 | 7.7 | 0.0 | 0.0 | Balanced |
| MT-I | 105.313567 | 0.3581 | 9.5 | 0.8 | 112.03203 | 1.08642562 | 9.0 | 1.3 | 0.5 | Arrestin |
| MT-II | 99.7405 | 0.17418 | 9.8 | 1.1 | 92.588748 | 0.76032628 | 9.1 | 1.4 | 0.3 | Arrestin |
| Bremelanotide | 95.9236831 | 0.26485 | 9.6 | 0.9 | 99.991565 | 2.00909281 | 8.7 | 1.0 | 0.1 | Arrestin |
| THIQ | 80.49922 | 2.89068 | 8.4 | -0.2 | 76.842593 | 6.16595002 | 8.1 | 0.4 | 0.6 | Arrestin |

**Table S5.** Calculation of ligand mediated bias factor with MC4R^T312A^ variant.

| Ligand | G protein (Gα<sub>s</sub>) |  |  |  | β-arrestin 2 |  |  |  | Receptor bias |  |
| --- | --- | --- | --- | --- | --- | --- | --- | --- | --- | --- |
|  | Emax (%)<sup>*</sup> | EC<sub>50</sub> (nM)<sup>#</sup> | Log (Emax/EC<sub>50</sub>) | ΔLog (Emax/EC<sub>50</sub>) | Emax (%)<sup>*</sup> | EC<sub>50</sub> (nM)<sup>#</sup> | Log (Emax/EC<sub>50</sub>) | ΔLog (Emax/EC<sub>50</sub>) | ΔΔLog (Emax/EC<sub>50</sub>) | Biased |
| α-MSH | 100 | 1.10408 | 9.0 | 0.0 | 100 | 8.01678063 | 8.1 | 0.0 | 0.0 | Balanced |
| MT-I | 113.514733 | 0.30832 | 9.6 | 0.6 | 100.94108 | 2.36047823 | 8.6 | 0.5 | -0.1 | G protein |
| MT-II | 116.467947 | 0.26122 | 9.6 | 0.7 | 93.335273 | 1.80301774 | 8.7 | 0.6 | -0.1 | G protein |
| Bremelanotide | 111.943929 | 0.29648 | 9.6 | 0.6 | 90.692695 | 4.13999675 | 8.3 | 0.2 | -0.4 | G protein |
| THIQ | 108.557827 | 2.09411 | 8.7 | -0.2 | 73.05212 | 33.2659553 | 7.3 | -0.8 | -0.5 | G protein |

**Table S6.** Calculation of ligand mediated bias factor with MC4R^S329A^ variant.

| Ligand | G protein (Gα<sub>s</sub>) |  |  |  | β-arrestin 2 |  |  |  | Receptor bias |  |
| --- | --- | --- | --- | --- | --- | --- | --- | --- | --- | --- |
|  | Emax (%)<sup>*</sup> | EC<sub>50</sub> (nM)<sup>#</sup> | Log (Emax/EC<sub>50</sub>) | ΔLog (Emax/EC<sub>50</sub>) | Emax (%)<sup>*</sup> | EC<sub>50</sub> (nM)<sup>#</sup> | Log (Emax/EC<sub>50</sub>) | ΔLog (Emax/EC<sub>50</sub>) | ΔΔLog (Emax/EC<sub>50</sub>) | Biased |
| α-MSH | 100 | 1.49279 | 8.8 | 0.0 | 100 | 5.17606832 | 8.3 | 0.0 | 0.0 | Balanced |
| MT-I | 111.896233 | 0.34198 | 9.5 | 0.7 | 95.672265 | 4.34510224 | 8.3 | 0.1 | -0.6 | G protein |
| MT-II | 123.4084 | 0.25177 | 9.7 | 0.9 | 90.51284 | 2.18776162 | 8.6 | 0.3 | -0.5 | G protein |
| Bremelanotide | 124.508874 | 0.34914 | 9.6 | 0.7 | 87.472403 | 5.99791076 | 8.2 | -0.1 | -0.8 | G protein |
| THIQ | 133.3145 | 2.24388 | 8.8 | -0.1 | 72.22116 | 43.9541615 | 7.2 | -1.1 | -1.0 | G protein |

**Table S7.** Calculation of ligand mediated bias factor with MC4R^S330A^ variant.

| Ligand | G protein (Gα<sub>s</sub>) |  |  |  | β-arrestin 2 |  |  |  | Receptor bias |  |
| --- | --- | --- | --- | --- | --- | --- | --- | --- | --- | --- |
|  | Emax (%)<sup>*</sup> | EC<sub>50</sub> (nM)<sup>#</sup> | Log (Emax/EC<sub>50</sub>) | ΔLog (Emax/EC<sub>50</sub>) | Emax (%)<sup>*</sup> | EC<sub>50</sub> (nM)<sup>#</sup> | Log (Emax/EC<sub>50</sub>) | ΔLog (Emax/EC<sub>50</sub>) | ΔΔLog (Emax/EC<sub>50</sub>) | Biased |
| α-MSH | 100 | 1.65577 | 8.8 | 0.0 | 100 | 10.5925373 | 8.0 | 0.0 | 0.0 | Balanced |
| MT-I | 105.942667 | 0.4121 | 9.4 | 0.6 | 98.248065 | 4.02717034 | 8.4 | 0.4 | -0.2 | G protein |
| MT-II | 102.423117 | 0.35727 | 9.5 | 0.7 | 93.020203 | 1.77010896 | 8.7 | 0.7 | 0.1 | Arrestin |
| Bremelanotide | 96.4665796 | 0.55208 | 9.2 | 0.5 | 88.895595 | 6.36795521 | 8.1 | 0.2 | -0.3 | G protein |
| THIQ | 93.43477 | 3.94457 | 8.4 | -0.4 | 72.415998 | 41.6869383 | 7.2 | -0.7 | -0.3 | G protein |

**Table S8.**
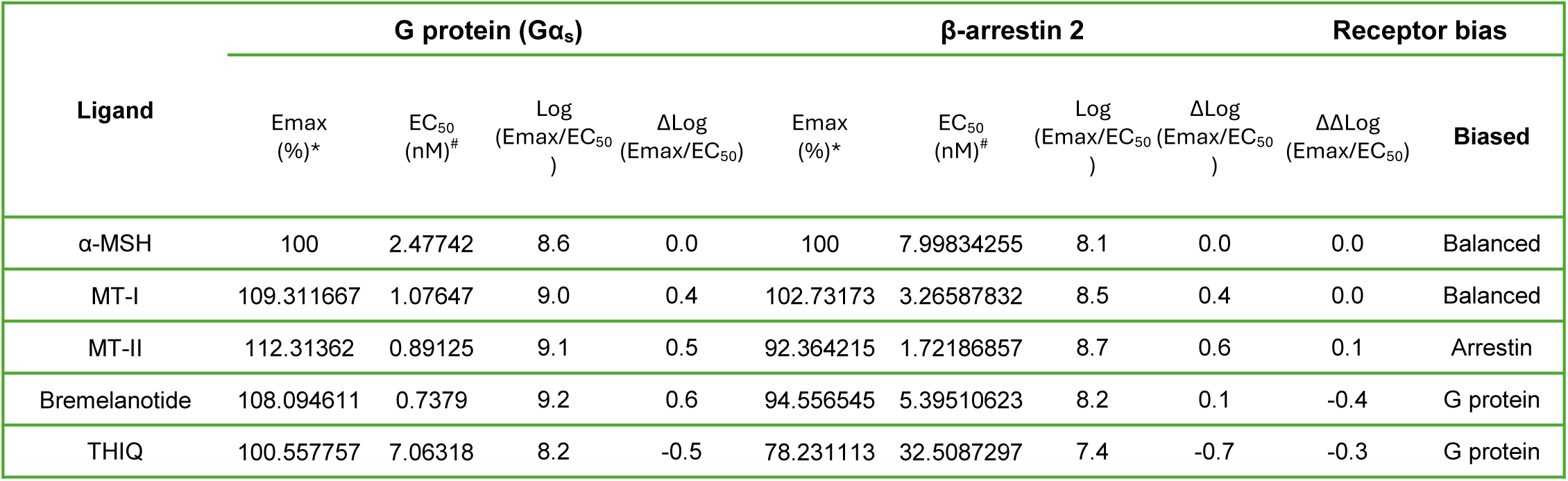
Calculation of ligand mediated bias factor with MC4R^S329A:S330A^ variant.

| Ligand | G protein (Gα<sub>s</sub>) |  |  |  | β-arrestin 2 |  |  |  | Receptor bias |  |
| --- | --- | --- | --- | --- | --- | --- | --- | --- | --- | --- |
|  | Emax (%)<sup>*</sup> | EC<sub>50</sub> (nM)<sup>#</sup> | Log (Emax/EC<sub>50</sub>) | ΔLog (Emax/EC<sub>50</sub>) | Emax (%)<sup>*</sup> | EC<sub>50</sub> (nM)<sup>#</sup> | Log (Emax/EC<sub>50</sub>) | ΔLog (Emax/EC<sub>50</sub>) | ΔΔLog (Emax/EC<sub>50</sub>) | Biased |
| α-MSH | 100 | 2.47742 | 8.6 | 0.0 | 100 | 7.99834255 | 8.1 | 0.0 | 0.0 | Balanced |
| MT-I | 109.311667 | 1.07647 | 9.0 | 0.4 | 102.73173 | 3.26587832 | 8.5 | 0.4 | 0.0 | Balanced |
| MT-II | 112.31362 | 0.89125 | 9.1 | 0.5 | 92.364215 | 1.72186857 | 8.7 | 0.6 | 0.1 | Arrestin |
| Bremelanotide | 108.094611 | 0.7379 | 9.2 | 0.6 | 94.556545 | 5.39510623 | 8.2 | 0.1 | -0.4 | G protein |
| THIQ | 100.557757 | 7.06318 | 8.2 | -0.5 | 78.231113 | 32.5087297 | 7.4 | -0.7 | -0.3 | G protein |

**Table S9.** Summary of α-MSH–induced signaling and trafficking profiles of MC4R and regulatory-site mutants. The table summarizes the relative responses of MC4R^WT^ and the indicated H8 and C-terminal phosphorylation-site mutants following stimulation with α-MSH. Responses are compared across receptor trafficking to endosomes, ERK1/2 activation, SRE-RE (Gα_s_), NFAT-RE (Gα_q/11_), and SRF-RE (Gα_12/13_) signaling pathways. Arrows indicate the relative magnitude of the observed response, with increasing numbers of arrows representing progressively stronger responses.

|  | H-8 Mutants |  |  | C-Terminus Mutants |  |  |
| --- | --- | --- | --- | --- | --- | --- |
|  | MC4R<sup>WT</sup> | MC4R<sup>S306A</sup> | MC4R<sup>T312A</sup> | MC4R<sup>S329A</sup> | MC4R<sup>S330A</sup> | MC4R<sup>S329A:S330A</sup> |
| cAMP Accumulation | ↑↑↑↑ | ↑↑ | ↑↑ | ↑↑ | ↑↑ | ↑↑ |
| β-arr2 Recruitment | ↑↑↑↑ | ↑ | ↑↑ | ↑↑↑↑ | ↑↑↑↑ | ↑↑↑↑ |
| Receptor Endocytosis | ↑↑↑↑ | ↑ | ↑↑ | ↑↑ | ↑↑↑↑ | ↑↑↑↑ |
| ERK 1/2 Activation (Early) | ↑↑↑↑ | ↑↑ | ↑↑ | ↑ | ↑ | ↑↑ |
| SRE Response (ERK) | ↑↑↑↑ | ↑ | ↑↑ | ↑↑ | ↑↑ | ↑↑ |
| NFAT Response (Gα<sub>q</sub>) | ↑↑↑↑ | ↑ | ↑ | ↑↑↑ | ↑ | ↑↑ |
| SRF-RE (Gα<sub>12/13</sub>) | ↑↑↑↑ | ↑ | ↑↑↑ | ↑ | ↑ | ↑ |

**Table S10.**
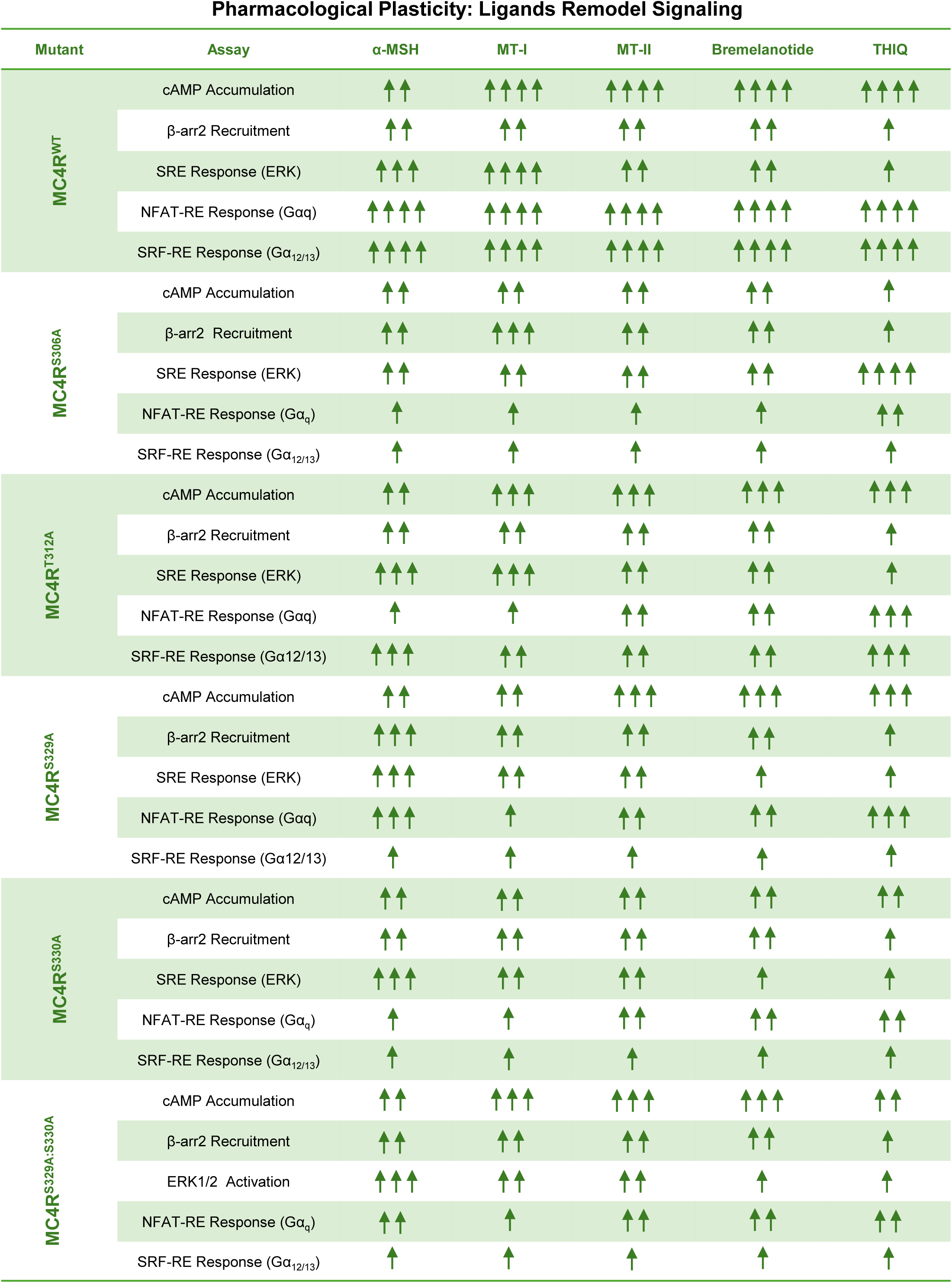
Summary of ligand-dependent signaling responses of MC4R and regulatory-site mutants: The table provides an integrated summary of the signaling responses of MC4R^WT^ and phosphorylation-site mutants (S306A, T312A, S329A, S330A, and S329A/S330A) following stimulation with the indicated ligands. Responses are summarized across cAMP accumulation, β-arrestin 2 recruitment, SRE based ERK activation, NFAT-RE (Gα_q/11_), and SRE-RE (Gα_12/13_) signaling assays. The summarized responses are derived from the experimental data presented in Figures 6, 8, and 9 and are consistent with the Emax values reported in Supplementary Tables S3–S8. Arrows denote the relative magnitude of the observed response, with an increasing number of arrows indicating progressively stronger responses.

